# Cargo-directed assembly of nonviral nucleocapsid with controlled size

**DOI:** 10.1101/2025.11.06.684264

**Authors:** Kenya Tajima, Yusuke Sakai, Naohiro Terasaka

## Abstract

Precise packaging of diverse cargo within self-assembling protein cages of defined size and shape is essential for many biotechnological applications, yet cellular expression offers limited control over loading. Here, we developed a system for *in vitro* cargo-directed reconstitution of a split, laboratory evolved nonviral nucleocapsid (spNC-4). Independently expressed and purified spNC- 4 capsid protein subunits were mixed and assembled with cargo molecules in a cooperative manner. As an authentic cargo, mRNA is packaged into a 30 nm-spheric nucleocapsid *in vitro*, closely matching to spNC-4 expressed in cells. In this system, a diverse range of cargo molecules, including cognate nucleocapsid mRNA, noncognate RNA, RNA complexed with positively supercharged fluorescent protein, and linear double-stranded DNA are encapsulated within the 30 nm-spheric nucleocapsids *in vitro*. Moreover, the packaging of 30 nm-spherical or rod-shaped DNA origamis as templates induce morphological alterations of the nucleocapsids, resulting in the formation of enlarged 60 nm-spherical structure or rod-shaped structure, respectively. This split-protein, cargo-dependent system provides versatile and programmable control over both composition and architecture of nonviral protein cages, creating a general platform for enzyme nanoreactors, targeted delivery, and vaccine development.

## Main text

Self-assembling protein cages are potential platforms for creating functional nanomaterials due to their monodisperse, hollow, and symmetrical architectures.^1–3^ They have been developed from viral capsids,^4,5^ through *de novo* protein design^6^ and by repurposing natural protein microcompartments.^7,8^ Encapsulation of various cargo molecules within protein cages has enabled functionalization, often driven by interactions between the cargo and the protein cage. While protein cages can be produced within cells, an alternative approach involves reassembling disassembled protein subunits *in vitro*, thereby expanding the scope of what can be encapsulated. Recent studies have demonstrated the successful packaging of DNA,^9–12^ scaffolded DNA origami,^13–16^ RNA,^17–19^ proteins,^20,21^ both nucleic acids and proteins,^22–26^ small molecules,^27^ metal nanoparticles^28^ and quantum dots.^29^

Among the available scaffolds, nonviral protein cages are of particular interest due to their high productivity, physical robustness, and inherent biosafety. In particular, *Aquifex aeolicus* lumazine synthase (AaLS) has been widely engineered due to its structural stability and functional versatility.^30^ For instance, AaLS cages have been engineered to encapsulate specific proteins via the peptide tag of its native cargo, riboflavin synthase,^31^ or to package positively charged proteins into negatively charged lumens through electrostatic interactions.^32–38^ This strategy has enabled AaLS cages to function as nanoreactors and protein delivery vehicles.^34–37,39^ Conversely, engineering a positively charged lumen enabled the nonspecific packaging of RNA, protecting it from degradation.^17,40,41^ To achieve specific RNA packaging, we previously introduced an RNA binding λN^+^ peptide into the lumen of circularly permutated AaLS and inserted BoxB stem-loop RNAs as packaging signal into its mRNA. Through multiple rounds of laboratory evolution, we generated an artificial virus-like nucleocapsid (NC-4) that packages its own mRNA in *Escherichia coli* cells.^17,42^ NC-4 forms a homogenous 240-subunit icosahedral capsid that packages, on average, 2.5 full-length mRNA molecules per capsid, along with nonspecifically packaged endogenous RNA. Notably, the absence of the λN^+^ peptide resulted in heterogeneous protein assemblies, suggesting that cargo RNA acts as template for uniform protein assembly and that NC-4 exhibits inherent morphological flexibility.

Harnessing this flexibility for controlled *in vitro* assembly, however, remains challenging. The NC-4 capsid protein spontaneously self-assembles into nucleocapsids during cellular expression. Furthermore, isolated NC-4 subunits are prone to aggregation and precipitation *in vitro.*^43^ Here, we overcome these limitations using a split-protein strategy, where the NC-4 capsid protein is divided into an N-terminal and a C-terminal fragments (N subunit and C subunit, respectively, Figure 1a). Inspired by their ability to co-assemble *in vivo,*^44^ we independently expressed and purified the N subunit and the C subunit. We demonstrate that these subunits can be controllably reassembled *in vitro* only in the presence of cargo, enabling the encapsulation of a wide array of molecules, including RNA, DNA, and proteins. Finally, we show that scaffolded DNA origami templates^45^ can direct the assembly process to produce different morphologies, yielding larger spherical or rod-shaped nanostructures. This work establishes a versatile platform based on split nonviral nucleocapsids, unlocking their potential for the programmable, bottom-up fabrication of functional nanomaterials.

**Figure 1:**
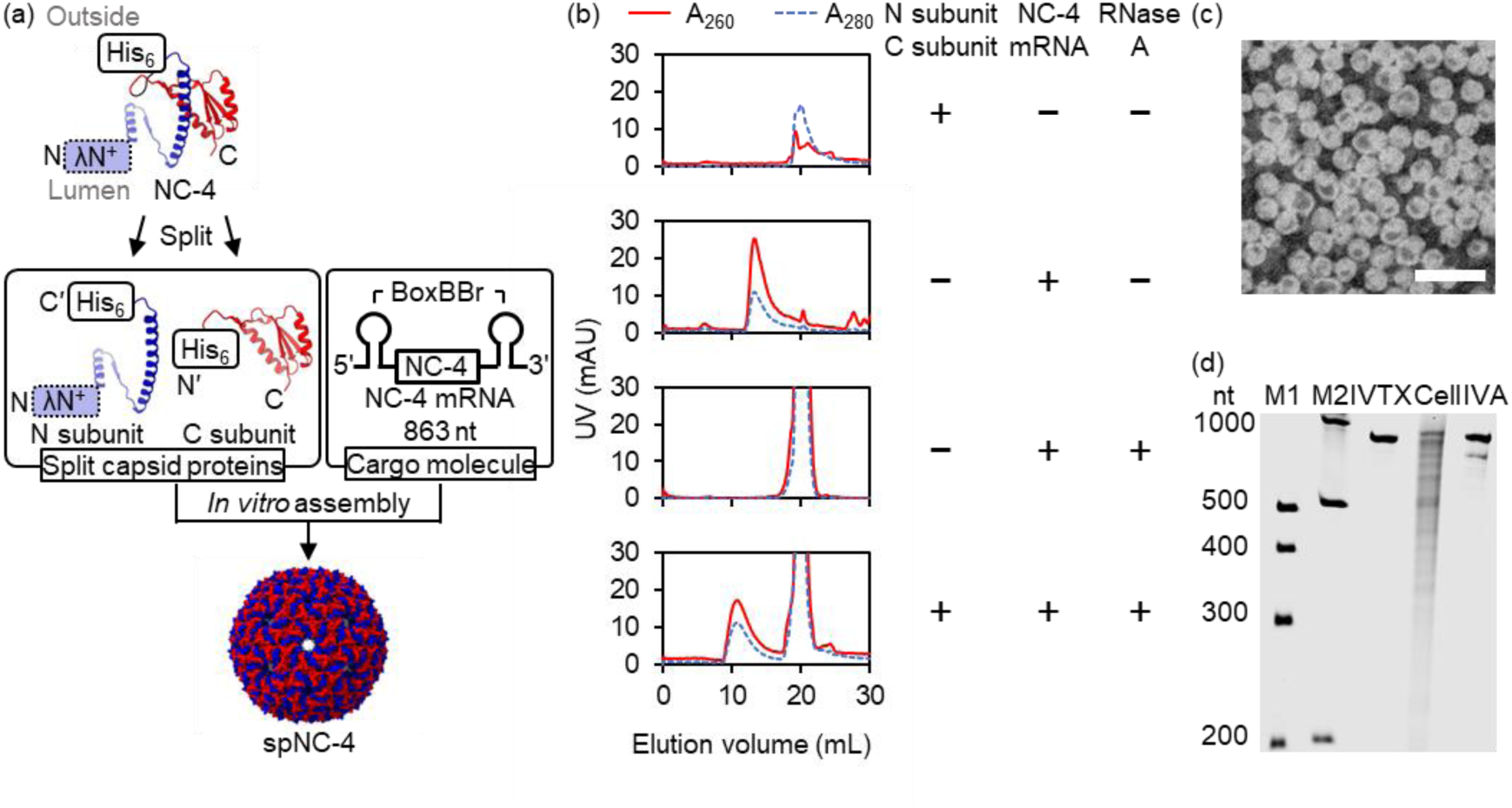
*In vitro* assembly of split capsid proteins and RNAs into a virus-like nucleocapsid. **(a)** *In vitro* assembly of split NC-4 (spNC-4) proteins and NC-4 mRNA. Split capsid proteins N subunit and C subunit, which have hexahistidine-tag for affinity purification, and *in vitro*- transcribed NC-4 mRNA were mixed *in vitro* to assemble into a capsid. λN^+^: A cationic peptide binding to an RNA motif BoxB. Molecular graphics were generated by using PDB ID 7A4J on UCSF ChimeraX. **(b)** Size-exclusion chromatographs of N subunit and C subunit, NC-4 mRNA, NC-4 mRNA degraded by RNase A and *in vitro* assembly of N subunit, C subunit and NC-4 mRNA challenged by RNase A. UV traces are shown with red lines and blue break lines for absorbance at 260 nm (A260) and 280 nm (A280), respectively. **(c)** A negative-stain transmission electron microscopic (TEM) images of spNC-4 prepared *in vitro* with NC-4 mRNA. Scale bar: 100 nm. (**d**) Denaturing polyacrylamide gel electrophoretic (PAGE) analysis of extracted RNA after *in vitro* assembly and RNase A challenge. M1 and M2: RNA markers, IVTX: *in vitro* transcribed NC-4 mRNA, Cell: RNA extracted from NC-4 expressed in *E. coli* after RNase A challenge, IVA: RNA extracted from spNC-4 assembled *in vitro* with N subunit, C subunit and NC-4 mRNA after RNase A challenge.

## Results

### RNA Cargo-Dependent *In Vitro* Assembly of a Split Artificial Nucleocapsid

To enable controlled *in vitro* assembly, the N subunit and the C subunit bearing hexahistidine tags of the split NC-4 (spNC-4) were independently expressed in *E. coli* and purified from inclusion bodies under denaturing conditions in the presence of urea (Figure S1a).^46^ During refolding attempts via buffer exchange to remove urea, the N subunit remained soluble, but the C subunit, which has a more hydrophobic surface, formed insoluble aggregates. To circumvent this, we developed a co-assembly strategy: the purified N subunit was first refolded, while the C subunit was kept denatured in urea. The two subunits were then mixed in the presence of cargo mRNA, and urea was rapidly diluted. This process triggered the simultaneous refolding of the C subunit and its cargo-directed assembly with the N subunit into nucleocapsids (Figure 1a). Unencapsulated RNA was subsequently removed by RNase A treatment.

Size-exclusion chromatography (SEC), monitored at 260 nm (A₂₆₀) and 280 nm (A₂₈₀) UV absorption, confirmed the formation of a stable complex, which appeared as a sharp peak at an elution volume of ∼11 mL, consistent with previously characterized *in vivo*-assembled NC-4 and spNC-4 nucleocapsids (Figure 1b),^17,44^ This peak, which was resistant to RNase A, was absent in control experiments lacking any of the three components (N subunit, C subunit, or RNA), confirming that assembly occurred in cargo-dependent manner. Negative-stain transmission electron microscopy (TEM) analysis of the peak fractions revealed monodisperse, spherical particles approximately 30 nm in diameter, matching the morphology of the native nucleocapsids (Figure 1c). Denaturing polyacrylamide gel electrophoresis (PAGE) of the RNA extracted from these particles confirmed that they protected the full-length mRNA cargo from nuclease (Figure 1d). These results demonstrate that the mRNA acts as a scaffold, recruiting the split protein fragments and templating their co-assembly into protective nanocontainers, a cooperative mechanism reminiscent of assembly in protease-induced NC-4 assembly^43^ and in many natural viruses.^47^

### Optimization of the Assembly Conditions

The assembly process relies on a balance of electrostatic interactions between RNA and the N subunit, and hydrophobic interactions between the protein subunits. To maximize the assembly yield, we systematically optimized the conditions by varying NaCl concentration, pH, and temperature. Using a fixed molar ratio of the N subunit, the C subunit, and NC-4 mRNA (130:130:1), we found that assembly was most efficient under physiological conditions (50 mM NaCl, pH 7.4, 37 °C), reaching completion within 30 minutes (Figure S1b–i). At these conditions, 28% of the proteins and 30% of the mRNAs were assembled into nucleocapsids (Figure S1j). Next, we titrated the NC-4 mRNA concentration from 0.23 μM to 3.72 μM. The maximum protein assembly yield (52%) was observed at 1.40 μM NC-4 mRNA; however, further increases in NC- 4 mRNA reduced the assembly yield (Figure S1j). Under conditions of 0.23 μM NC-4 mRNA, an average of 1707 nucleotides were packaged per nucleocapsid (Figure S1k), corresponding to about two copies of NC-4 mRNA. This was comparable to the ∼2500 nucleotides packaged by NC-4 expressed in *E. coli*^17^ and approximately half of the ∼3400 nucleotides packaged by spNC- 4 expressed in *E. coli*.^44^ For subsequent experiments, we used 0.23 μM NC-4 mRNA, which provided a robust yield and cargo loading.

### *In vitro* packaging of non-cognate RNAs

We next explored the system’s packaging versatility. NC-4 and spNC-4 expressed in *E. coli* contain endogenous RNAs in addition to the mRNAs carrying packaging signals.^17^ We designed three RNAs derived from NC-4 mRNA: a 60-nt BoxB-Broccoli (BoxBBr) packaging signal RNA, as well as 145-nt and 445-nt 3′-truncated NC-4 mRNAs (Figure S2a-c, top). Because NC-4 mRNA includes packaging signal-like structures in its open reading frame, the 145-nt and 445-nt truncated NC-4 mRNAs carry three and four packaging signals, respectively.^17^ All three RNAs stimulated nucleocapsid formation, as confirmed by negative-stain TEM analysis (Figure S2a-c, middle). SEC analysis further showed that longer RNAs and/or a greater number of packaging signals improved nucleocapsid yield and RNA packaging (Figure S2a-c, bottom, Figure S2d, e). Furthermore, nucleocapsid formation was also observed with non-cognate RNAs, such as super-folding GFP (sfGFP) mRNA with or without two BoxBBr packaging signals, as confirmed by negative-stain TEM analysis (Figure S2f, g, middle). We found that sfGFP mRNA lacking packaging signals showed a nucleocapsid yield (28%) comparable to that of NC-4 mRNA, whereas sfGFP mRNA containing two BoxBBr tags exhibited a higher yield (38%) and enhanced RNA packaging compared to NC-4 mRNA (Figure S2f, g, bottom, Figure S2h, i).

### *In vitro* co-packaging of RNA and positively supercharged GFP

In earlier studies, AaLS variants with negatively charged lumens successfully encapsulated positively charged proteins via electrostatic interactions.^32–37^ Inspired by these findings and by the successful packaging of negatively charged RNA lacking the BoxBBr sequence into spNC-4, we attempted *in vitro* packaging of negatively charged GFP(−30)^48^ into spNC-4. However, protein cage formation was not observed (Figure S3a).

We therefore attempted to co-package a supercharged protein, sfGFP(+4), GFP(+36) and GFP(−30), along with the mRNA cargo. SEC analysis revealed A₂₈₀ and A₂₆₀ peaks at ∼11 mL in all three conditions; however, a GFP fluorescence peak appeared only when 1.25 μM of positively charged GFP(+36) was present (Figure S3b). This observation likely reflects electrostatic interactions between RNA and GFP(+36). At GFP(+36) concentrations higher than 1.25 μM, precipitation occurred, resulting in no detectable UV absorbance or GFP fluorescence (Figure S3b, right panels). Negative-stain TEM analysis confirmed nucleocapsid formation in the presence of 1.25 μM GFP(+36) (Figure 2a). When spNC-4 assembled *in vivo* was mixed with GFP(+36) *in vitro* and analyzed by SEC, GFP(+36) fluorescence appeared in all eluted fractions, suggesting that GFP(+36) binds to the negatively charged outer surface of spNC-4 (Figure 2b, Figure S3c). Because GFP(+36) (28.5 kDa) is larger than RNase A (13.7 kDa), it cannot diffuse into the lumen of assembled spNC-4.^17,44^ To confirm true encapsulation versus nonspecific surface binding, we developed a high-salt wash (1 M NaCl) capable of stripping surface-adsorbed proteins (Figure 2b, second top). Even after this stringent wash, a significant GFP(+36) fluorescence signal remained co-located with the nucleocapsid peak in SEC, and intact capsids were confirmed by TEM (Figure 2b, c), demonstrating that both NC-4 mRNA and GFP(+36) were successfully co-packaged. The number of encapsulated GFP(+36) molecules was tunable, reaching approximately six molecules per capsid at 1.25 μM GFP(+36) (Figure 2d). Assembly yields and packaged RNA nucleotides (approximately 2000 nt per a nucleocapsid) were comparable to those observed for RNA-only packaging (Figure S3d, e), with a yield of about 10% based on protein concentration. This result demonstrates the system’s ability to co-package distinct classes of macromolecules—nucleic acids and proteins—within a single nanoparticle.

**Figure 2:**
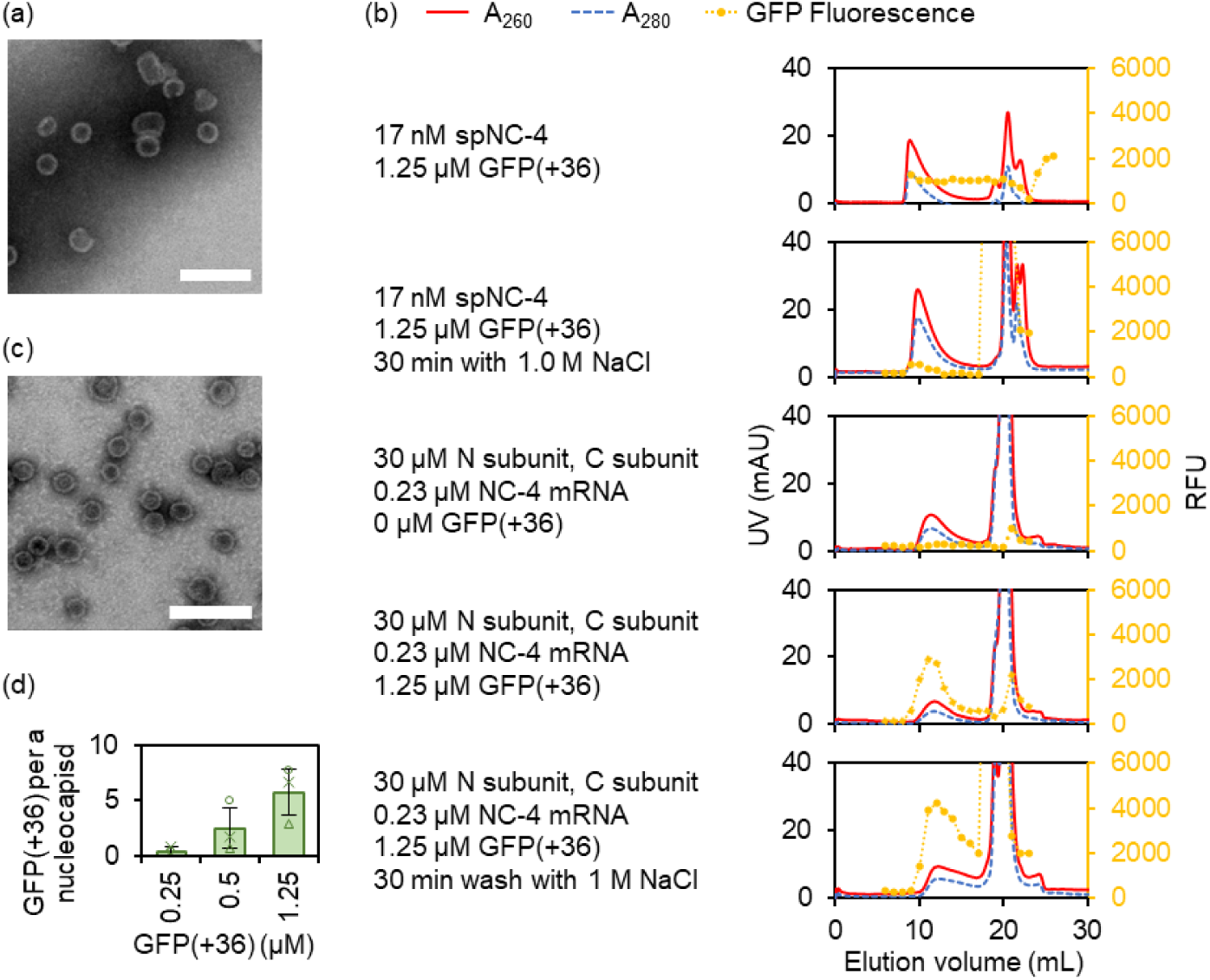
*In vitro* assembly of split capsid proteins, RNA and positively-supercharged protein. **(a), (c)** Negative-stain TEM images of spNC-4 prepared *in vitro* with N subunit, C subunit, NC- 4 mRNA and 1.25 μM of GFP(+36) (a) without or (c) after incubation with 1.0 M NaCl for 30 minutes. Scale bars: 100 nm. **(b)**SEC analysis showing GFP(+36) packaged into spNC-4 lumen. Preassembled spNC-4 mixed with 1.25 μM of GFP(+36) without NaCl incubation (top) or incubated with 1 M NaCl for 30 minutes (second top), *in vitro* assembly of 30 μM each N subunit and C subunit and 0.23 μM NC-4 mRNA without GFP(+36) (middle), with 1.25 μM of GFP(+36) without NaCl incubation (second bottom), and with 1.25 μM of GFP(+36) incubated with 1 M NaCl for 30 minutes (bottom). RFU: relative fluorescence unit. **(d)** Number of packaged GFP(+36) molecules per a nucleocapsid assembled with 0, 0.25, 0.5, or 1.25 μM GFP(+36) with 1.0 M NaCl washing. The data is mean ± s.d. for three independent replicates tested.

### DNA Origami as a Template for Morphological Control

Finally, we investigated whether DNA could template assembly. All DNA samples were treated with RNase A to eliminate assembly driven by any contaminating RNA. First, the N subunit, the C subunit, and salmon sperm dsDNA (≤ 2000 bp) were mixed at different protein-to-DNA ratios, and the resulting nucleocapsids were analyzed by 0.6% agarose gel electrophoresis (AGE) (Figure S4). When the dsDNA concentration was fixed at 100 ng/μL, faint DNA bands appeared with 30 or 60 μM each of the N subunit and the C subunit, overlapping with Coomassie-stained protein bands (Figure S4, lane 7 and 8). The electrophoretic mobility of these bands resembled that of the authentic NC-4 nucleocapsid assembled in *E. coli* (Figure S4, lane 1). Negative-stain TEM analysis revealed partial nucleocapsid formation, indicating that linear dsDNA promoted only limited assembly. (Figure 3a). When the concentration of the N subunit and the C subunit was kept at 30 μM each, DNA bands overlapping with protein bands were also observed at dsDNA concentrations of 2500, 1200, or 625 ng/μL (Figure S4, lane 10, 11 and 12). Next, we investigated whether nucleocapsid assembly confers protection against nuclease digestion. Naked dsDNA and dsDNA packaged within spNC-4 were incubated with DNase I for up to six hours (Figure 3b). Whereas naked salmon dsDNA was fully degraded within 15 minutes (Figure 3b, lane 2 to 7), DNA bands co-migrating with the protein bands persisted even after six hours of DNase I treatment in the presence of N subunit and C subunit (Figure 3b lane 8 to 13) DNA bands co-migrating with the protein bands persisted even after six hours of DNase I treatment in the presence of N subunit and C subunit.

**Figure 3:**
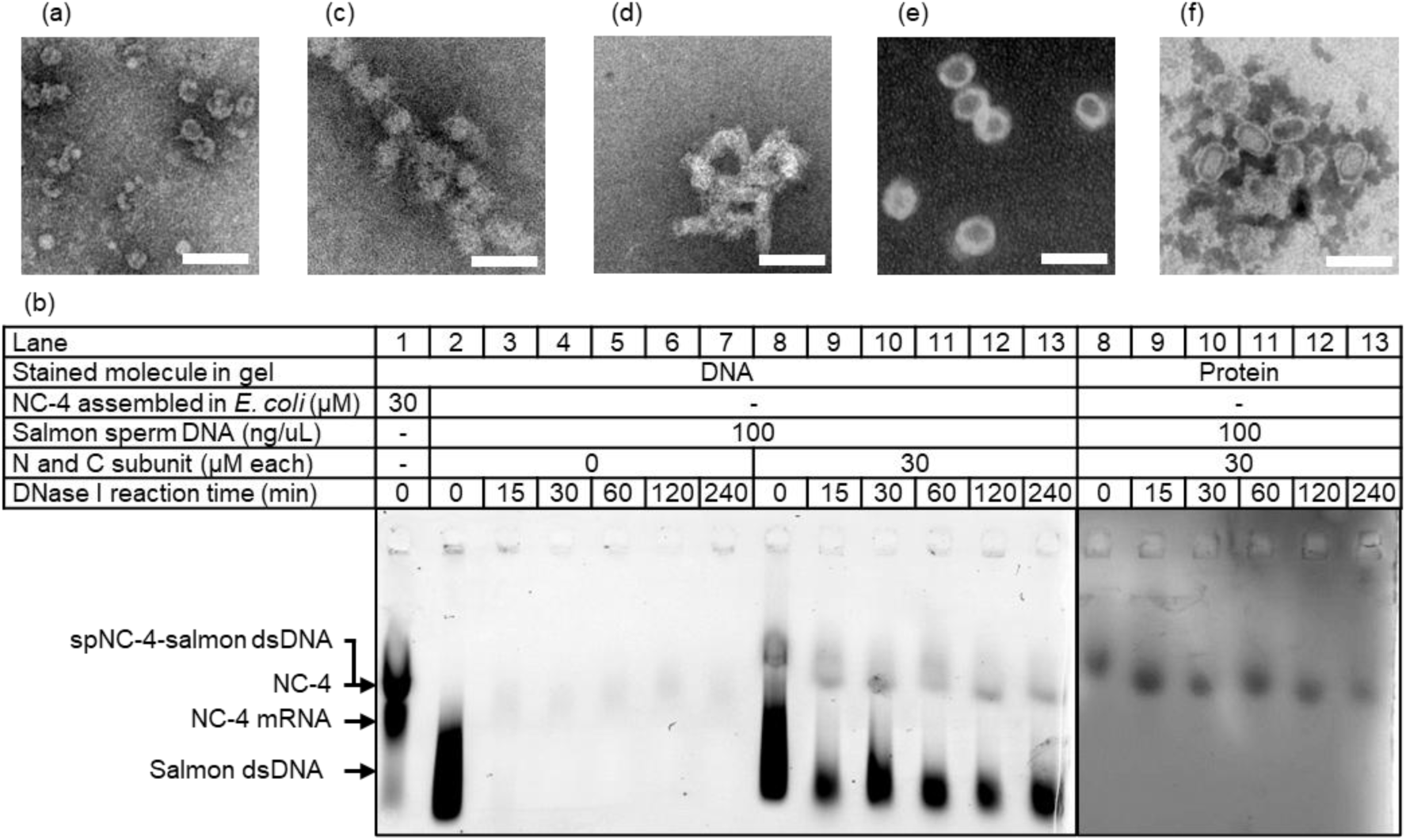
*In vitro* assembly of split capsid proteins and linear dsDNA or structured DNA origamis. **(a)** A negative-stain TEM image of nucleocapsids prepared with 30 μM each N subunit, C subunit and 100 ng/μL salmon sperm dsDNA. **(b)** DNase I challenge nucleocapsids encapsulating salmon sperm dsDNA*. In vitro* assembly reaction mixture was challenged by DNase I for 0, 15, 30, 60, 120 or 140 minutes and separated by 0.6% agarose gel electrophoresis (AGE). **(c), (e)** A negative-stain TEM image of 2 nM DNA origami sphere (c) alone with 20 mM MgCl2 and (e) assembled with 24 μM each N subunit and C subunit with 12 mM MgCl2. **(d), (f)** A negative-stain TEM image of a DNA origami rod (d) alone and (f) 1 nM DNA origami rod assembled with 6 μM each N subunit and C subunit with 20 mM MgCl2. Scale bars: 100 nm.

We sought to package scaffolded DNA origami cargos into the nucleocapsid to control its overall morphology. Previous studies have shown that encapsulating a structured cargo can direct assembly patterns and thereby influence protein cage morphology.^13,16,49,50^ DNA origami technology offers nanoarchitectures with user-defined three-dimensional shapes and sizes,^45^ making it an ideal morphological template for nucleocapsid assembly. We designed two types of DNA origami architectures, a sphere (30 nm in diameter) and a rod (60 nm in height and 25 nm in diameter), to generate structures distinct from the ∼30 nm spherical form of NC-4 and spNC-4.^17,43,44^ Staple strands and scaffold strands were mixed at a 1:10 ratio and annealed overnight using a thermal ramp in a buffer containing optimal Mg²⁺ concentrations. Negative-stain TEM analysis confirmed the successful assembly of the DNA origami cores (Figure 3c, d). When the DNA origami sphere was packaged *in vitro*, negative-stain TEM revealed spherical particles ∼60 nm in diameter, approximately twice the size of NC-4 and spNC-4 (Figure 3e). In contrast, packaging the rod-shaped DNA origami yielded rectangular structures ∼63 nm in height and ∼38 nm in width (Figure 3f). These results demonstrate that DNA origami cargos can modulate the structural morphology of the non-viral nucleocapsid.

## Discussion

In this study, we established a robust *in vitro* reconstitution system for a laboratory-evolved nucleocapsid, demonstrating its remarkable versatility in packaging a wide array of cargo molecules including RNA, proteins, and DNA. To obtain the capsid protein components necessary for *in vitro* reconstitution, folded N subunit and denatured C subunit of split NC-4 were independently prepared because the disassembled NC-4 capsid protein was poorly recovered from

*E. coli.*^43^ Subsequently, *in situ* refolding and cooperative assembly with cargo molecules was achieved. Compared to the previous study that demonstrated *in vitro* assembly of NC-4 triggered by protein cleavage,^43^ our approach simply mixing protein and cargo molecules resulted in a higher yield of nucleocapsid (50% of proteins were assembled) in shorter assembly time (30 minutes to about 2 days). Additionally, recent studies have reported that reassembly of an artificial protein nanocage through the mixing of purified split protein units.^51^ These studies demonstrates that split/denature/refolding/assembly strategy could be broadly applicable to other protein cages whose disassembled proteins were poorly isolated. Furthermore, this approach can facilitate the modification of capsids via newly generated protein termini.^44^

The *in vitro* packaging of RNA with various length was demonstrated. Each nucleocapsid particle encapsulated 1.9 to 2.5 molecules of full-length mRNA and protected cargo from RNase A degradation, exhibiting a morphology nearly identical to that of NC-4 and spNC-4 expressed in cell.^17,44^ Nucleocapsid formation was enhanced at lower salt concentration likely due to the stronger affinity of λN^+^ peptide for the BoxB RNA packaging signal under low salt concentration.^52^ This preference for low salt concentration suggested an *en masse* assembly mechanism, in which capsid proteins initially bound to RNA in a disordered manner and subsequently undergo cooperatively assembly into ordered nucleocapsids.^53^ Regardless of RNA length and secondary structure, no apparent morphological changes in nucleocapsid were observed. However, nucleocapsid assembly was more efficient in the presence of longer RNAs. The net negative charge of packaged long RNAs (−1730, −1974 and −2030 for two copies of NC- 4 mRNA, GFP mRNA and GFP mRNA with two BoxBBr packaging signals) exceeded net positive charge of inner surface of spNC-4 (+1440 as a 240-mer complex). This observation is consistent with the overcharged nature of single-stranded RNA (ssRNA) genomes^54^ and the preferential packaging of longer RNAs in certain ssRNA viruses.^55^ The *in vitro* packaging of GFP mRNA into spNC-4 was not significantly affected by the presence of packaging signals. In some natural viruses, such as bacteriophage MS2 and satellite tobacco necrosis virus (STNV), packaging signals play an essential role in assembly,^56^ facilitating a conformational switch in the capsid proteins upon binding.^57,58^ This suggests that nonspecific electrostatic interactions between RNA and multiple N subunit proteins are sufficient to drive nucleocapsid assembly without conformational switch. This *in vitro* study suggests that packaging signals presumably contribute to selective packaging of own mRNA via the interaction between BoxB and λN^+^ peptides of NC- 4 *in vivo*, which is consistent to the results of cellular expression of nucleocapsids from mRNA without BoxBBr signals.^42^

Positively-charged GFP(+36) was co-packaged with NC-4 mRNA into spNC-4 via electrostatic interactions with NC-4 mRNA. In our assembly system, protein cage assembly was not stimulated by negatively-charged GFP(−30) in the absence of NC-4 mRNA. In the case of RNA packaging, as discussed above, capsid proteins are bridged by RNA of sufficient length before nucleation of nucleocapsids is initiated^43^. We speculated that GFP(−30) was too small to bridge λN^+^ peptides of N subunit to initiate the nucleation. In this study, co-packaging was achieved by simply mixing two purified capsid proteins with a cargo protein and long mRNA. This straightforward system would enable the co-packaging of various proteins and RNA by fusing GFP(+36) to proteins of interest. However, the number of packaged proteins was relatively low compared to the other strategies, such as the conjugation of nucleic acids to cargo proteins^23–25^. This limitation is likely due to weak non-specific electrostatic interactions between the cargo protein and RNA. Improving these interactions, for example, by introducing a natural RNA-binding protein^59^ or an RNA aptamer^60^ could enhance protein packaging efficiency, making this approach more compatible with cellular co-packaging systems.

In addition to RNA and protein, various types of DNA such as linear double strand DNA, spherical and rod-shaped DNA origami were successfully utilized as cargo molecules in this study. When packaging scaffolded DNA origami larger than original spNC-4, the assembly pattern of capsid proteins appeared to be reorganized. Such structural reorganization was also observed in NC-4 lacking λN^+^ peptide which exhibited heterogenous structures.^17^ As expected, the capsid proteins adopted their assembly patterns to cover the DNA origami surfaces, forming a 60-nm- diamter spherical structure or a 63-nm-high and 38-nm-wide rod-shaped structure. Although the detailed structure has not yet determined, the Caspar-Klug triangulation number^61,62^ for the 60- nm-diamter nucleocapsid packaging spherical DNA origami would be 16 (960 mer protein) if the trimeric quasi-symmetric unit was maintained. These results highlight the versatile packaging capability and structural flexibility of artificially evolved nonviral nucleocapsids.

In conclusion, our work establishes a versatile platform for the cargo-directed assembly of a split, nonviral nucleocapsid. This system not only encapsulates and protects a diverse range of functional molecules but also allows its morphology to be programmed using structured nucleic acid templates. This combination of packaging versatility and structural control provides a powerful toolkit for the future development of enzyme nanoreactors, targeted delivery vehicles, and novel vaccine platforms.

## Author contributions

N.T. supervised this study. K.T. carried out *in vitro* nucleocapsid assembly and characterization.

Y.S. designed and prepared DNA origami architectures. All the authors participated in manuscript preparation.

## Methods

### Chemicals, DNA oligos and synthetic gene fragments

All the chemicals without mention were purchased from Nacalai Tesque. Oligo DNAs and synthetic gene fragments were purchased from IDT. The M13mp18 phage genome (p7249), as well as its variant p8064, used as the scaffold strand for DNA origami rods and DNA origami spheres, respectively, were purchased from Tilibit Nanosystems.

### *In vitro* transcription and purification of RNAs

Full-length and truncated NC-4 messenger RNAs (mRNAs) were prepared by runoff *in vitro* transcription. DNA templates were prepared by PCR with corresponding primer pairs from plasmid pMG-dB-NC-4 using KOD One PCR master mix (TOYOBO). PCR-amplified templates were purified by FastGene Gel/PCR Extraction Kit (NIPPON Genetics). *In vitro* transcription reactions were performed using in-house expressed T7 RNA polymerase in reaction mixture (40 mM Tris-HCl pH = 8.0, 1 mM spermidine, 0.01%v/v Triton-X100, 10 mM DTT, 30 mM MgCl2, 5 mM each ATP, UTP, CTP (Combi-Blocks) and GTP (FUJIFILM), 30 mM KOH, 0.12 μM T7 RNA polymerase) at 37 ℃ for 3-18 hours. Template DNA was digested by adding RQ1 DNase (Promega) with DNase reaction buffer (40 mM Tris-HCl at pH = 8.0, 10 mM NaCl, 6 mM MgCl2, 1 mM CaCl2 as final concentration) at 37 ℃ for 1 hour. RNA was precipitated with isopropanol. RNA sample was purified by denaturing polyacrylamide gel electrophoresis (PAGE). Preparative denaturing PAGE gels were prepared in Tris/borate/EDTA (TBE) buffer supplemented with 8 M urea and 5% polyacrylamide. Polymerization was initiated using TEMED (7.5 μL per 10 mL gel solution) and 10% APS (10% in water, 100 μL per 10 mL gel solution). RNA bands were visualized by UV shadowing and excised with a scalpel. The gel pieces were crushed with a pipette tip and the RNA was extracted in water containing 0.3 M NaCl for 3-18 hours at room temperature. The RNA was precipitated by ethanol and the pellet was dissolved in water. The RNA concentration was quantified by UV absorbance at 260 nm by NanoDrop One^C^ (Thermo Fisher Scientific).

### Expression and purification of N subunit, C subunit, sfGFP(+4), GFP(**−**30), and GFP(+36)

All proteins were expressed in *E. coli* BL21(DE3)-gold cells (Agilent Technologies). Two-litter Erlenmeyer flasks containing 1000 mL LB medium with 100 mg/mL ampicillin were inoculated with 20 mL overnight cultures and incubated at 37 ℃ and at 200 rpm until OD600 reached 0.5. Protein expression was induced by adding IPTG (M&S TechnoSystems) to a final concentration of 0.3 mM. Cells were cultured at 20 ℃ for 16 hours and then harvested by centrifugation at 5000 *g* and 4 ℃ for 10 minutes. The cell pellet from one 1000-mL culture was resuspended in 40 mL LB medium, transferred to a 50-mL centrifugal tube. The medium used for transfer removed by centrifugation at 5000 *g* and 4 ℃ for 10 minutes, decantated, and aliquots of the cell pellets were frozen in liquid nitrogen and stored at −80 ℃ until purification.

For purification of N subunit and C subunit, a cell pellet corresponding to 1000 mL of culture volume was resuspended in 10 mL of buffer (50 mM Tris-HCl, 1 mM EDTA, 100 mM NaCl at pH 8.0) and lysed by sonication (60 cycles of 1 sec on and 1 sec off with 40% amplitude, Q500 Sonicator, QSONICA). The insoluble fraction in the lysate was precipitated by centrifugation with 12000 *g* at 4 ℃ for 30 minutes. The precipitation was resuspended in 10 mL buffer (50 mM Tris-HCl, 10 mM EDTA, 0.1 M NaCl, 0.5% Triton-X100 at pH 8.0) at 4 ℃ for 3 hours to resolubilize the cell membrane. Inclusion body of a protein was precipitated by centrifugation at 17000 *g* and 4 ℃ for 30 minutes and removing the supernatant and the inclusion body was stored at −80 ℃ until denaturing. The inclusion body extracted from 1000 mL of *E. coli* culture was resuspended in 10 mL of denaturing buffer (20 mM Tris-HCl, 8M urea at pH = 8.0) at 4 ℃ for 30 minutes and supernatant containing denatured N subunit or C subunit was recovered after centrifugation at 17000 *g* at 4 ℃ for 30 minutes. After addition of 100 mM of NaCl as final concentration, the supernatant was loaded onto 1 mL slurry of Ni(II)-NTA resin (Nuvia IMAC resin, Bio-Rad) in a gravity flow column. After incubation for 30 minutes, the resin was washed with 10 mL of buffer (50 mM Tris-HCl, 1 M NaCl, 20 mM imidazole without urea for N subunit, and that with 8 M urea for C subunit) and eluted with 4 aliquots of 1 mL of buffer (50 mM Tris-HCl, 300 mM NaCl, 300 mM imidazole without urea for N subunit, and that with 8 M urea for C subunit). The protein quantity was determined by UV absorbance at 280 nm by NanoDrop One^C^ (Thermo Fisher Scientific). The eluted fractions were frozen by liquid nitrogen and stored at −80 ℃.

For purification of sfGFP(+4), GFP(−30) and GFP(+36), a cell pellet corresponding to 1000 mL of culture volume was resuspended in 10 mL of native lysis buffer (50 mM sodium phosphate, 1 M NaCl, 20 mM imidazole at pH 7.4) and lysed by sonication (60 cycles of 1 sec on and 1 sec off with 40% amplitude, Q500 Sonicator, QSONICA). The soluble fraction in the lysate was recovered after centrifugation at 12000 *g* and 4 ℃ for 10 minutes and loaded onto 1 mL slurry of Ni(II)-NTA resin (Nuvia IMAC resin, Bio-Rad) in a gravity flow column. After incubation for 30 minutes, the resin was washed with 10 mL of native lysis buffer and eluted with 5 aliquots of 1 mL of native elution buffer (50 mM sodium phosphate, 300 mM NaCl, 100, 200, 300, 400 or 500 mM imidazole at pH 7.4). The eluted fractions were combined and dialyzed in 1 L of native lysis buffer without imidazole at 4 ℃ for at least 3 hours twice. The protein quantity was determined by using Pierce 660-nm Protein Assay Reagent (Thermo Fisher Scientific) with NanoDrop One^C^ (Thermo Fisher Scientific). Final solution was frozen by liquid nitrogen and stored at −80 ℃.

### Preparation of DNA origamis

The scaffold strand and staple strands are typically mixed at a 1:10 ratio in TAE buffer containing 10 mM MgCl₂ (rod-shaped core) or 20 mM MgCl₂ (sphere-shaped core). The mixture is annealed using a thermal cycler configured with the following multi-step thermal ramp: 95°C for 5 minutes, 95–66°C at a rate of -1°C per 8 minutes, 65–41°C at -1°C per 25 minutes, and 40– 25°C at -1°C per 8 minutes. Successful assembly was verified by agarose gel electrophoresis and negative-stained TEM.

### *In vitro* assembly of N subunit, C subunit and RNAs

Buffers containing appropriate concentration of NaCl and whose pH were adjusted, RNA, N subunit and C subunit were mixed in this order and solutions were mixed adequately by every addition. Reaction mixture was incubated at 37 ℃ for 30 minutes to 16 hours.

### *In vitro* assembly of N subunit, C subunit, NC-4 mRNA, and GFPs

Buffers containing appropriate concentration of NaCl and whose pH were adjusted, RNA, N subunit, GFPs and C subunit were mixed in this order and solutions were mixed adequately by every addition. Reaction mixture was incubated at 37 ℃ for 30 minutes to 16 hours. To wash GFP(+36) binding on the surface of nucleocapsid, NaCl was added to the reaction mixture with 1 M as final concentration and incubated at 37 ℃ for 30 minutes.

### Negative-stain transmission electron microscopy (TEM)

*In vitro* assembly reaction mixture was diluted by the same buffer used for assembly to make N subunit and C subunit concentration 6 μM. TEM grids (#783118953, JEOL or #6515, Nisshin-EM) were negatively glow discharged by soft mode with PIB-10 (Vacuum Device) for 30 sec before use. The solution was directly used for blotting on TEM grids. Grids were incubated with the nucleocapsid solution for 30 sec, washed twice and incubated for 15 sec with TEM staining solution (2% wt/vol aqueous phosphotungstate acid (TTAB Laboratories Equipment), pH 7.6, filtrated), dried and imaged by using H-7650 Zero.A transmission electron microscope (HITACHI).

### Size-exclusion chromatographic (SEC) quantification of nucleocapsids

For SEC analysis, RNase A (Thermo Fisher Scientific) was added into assembly reaction mixture with 10 μg/mL as final concentration after assembly reaction and incubated at 37 ℃ for 1 hour. The reaction mixture was filtered by a 0.45 μm filter unit (SLHVR04NL, Merck) and injected to NGC system (Bio-Rad) equipped with Superose 6 Increase 10/300 GL SEC column (Cytiva). The column was preequilibrated at room temperature by 200 mM NaCl SEC buffer (50 mM sodium phosphate, 200 mM NaCl, 5 mM EDTA at pH 7.4) or 1 M NaCl SEC buffer (50 mM sodium phosphate, 1 M NaCl, 5 mM EDTA at pH 7.4) only for co-packaging of NC-4 mRNA and GFPs. UV absorbance at 260 nm and 280 nm were measured by spectrometer built in NGC system and GFP fluorescence (excited by 480 nm laser and emission detected at 507 nm) was measured by EnSpire (PerkinElmer).

### RNA extraction from nucleocapsids assembled *in vitro*

RNA was extracted from spNC-4 nucleocapsid assembled *in vitro* after SEC purification. MgCl2 and RNase A (Thermo Fisher Scientific) were added to the nucleocapsid fraction with 5 mM and 10 μg/mL as final concentration and incubated at 37℃ for 1 hour. 5 equivalent volumes of TRIzol Reagent (Thermo Fisher Scientific) were added and RNA was extracted following the manufacturer’s instructions. Extracted RNA was separated by 8% denaturing PAGE with RNA markers (DM152, DM160, BioDynamics Laboratory) and *in vitro* transcribed NC-4 mRNA. Gel was stained by EzPreStain DNA&RNA (ATTO) and imaged by Typhoon FLA 9500 (Cytiva).

### *In vitro* assembly of N subunit, C subunit and salmon dsDNA or DNA origamis and separation by agarose gel electrophoresis

For *in vitro* assembly mediated by DNA, DNA was preincubated with 10 μg/mL of RNase A (Thermo Fisher Scientific) at 37 ℃ for 1 hour. 1x TAE buffer (4 mM Tris, 2 mM AcOH, 1 mM EDTA), appropriate concentration of MgCl2, N subunit, DNA and C subunit were mixed in this order and solutions were mixed adequately by every addition. Reaction mixture was incubated at 37 ℃ for 16 hours. Naked DNA was degraded by DNase I (Promega) for 15 minutes to 6 hours. DNase I reaction was quenched by 40 mM EDTA as final concentration. Reaction mixture was separated by 0.6% agarose gel electrophoresis without MgCl2 for salmon dsDNA prestained by EzPrestain DNA&RNA (ATTO) and imaged by Typhoon FLA 9500 (Cytiva). Proteins were stained by EzStain AQua (ATTO) and imaged by Printgraph CMOS I (ATTO).

### Molecular graphics and coulombic potential calculation

Molecular graphics and coulombic potential calculation were performed with UCSF ChimeraX v18.1^63^. Molecular graphics of NC-4 and sfGFP(+4) were modeled by utilizing 7A4J and 2B3P PDB files, respectively. Molecular graphics of GFP(−30) and GFP(+36) were generated from sfGFP(+4) structure on ChimeraX.

## Acknowledgement

We thank Materials Analysis Division, Core Facility Center, Research Infrastructure Management Center, Institute of Science Tokyo and “Advanced Research Infrastructure for Materials and Nanotechnology in Japan (ARIM)” of the Ministry of Education, Culture, Sports, Science and Technology (MEXT), Grant Number JPMXP1223UT0270 for use of TEM equipment. Molecular graphics and analyses performed with UCSF ChimeraX, developed by the Resource for Biocomputing, Visualization, and Informatics at the University of California, San Francisco, with support from National Institutes of Health R01-GM129325 and the Office of Cyber Infrastructure and Computational Biology, National Institute of Allergy and Infectious Diseases.

## Supporting Information Available

`●` SEC chromatographs of condition screening of NC-4 mRNA packaging.
`●` TEM images and SEC chromatographs of non-cognate RNA packaging.
`●` Gel images of salmon sperm DNA packaging.
`●` Uncropped gel images and SEC chromatographs.
`●` DNA, RNA and protein sequences used in this article.

---

**Figure S1.**
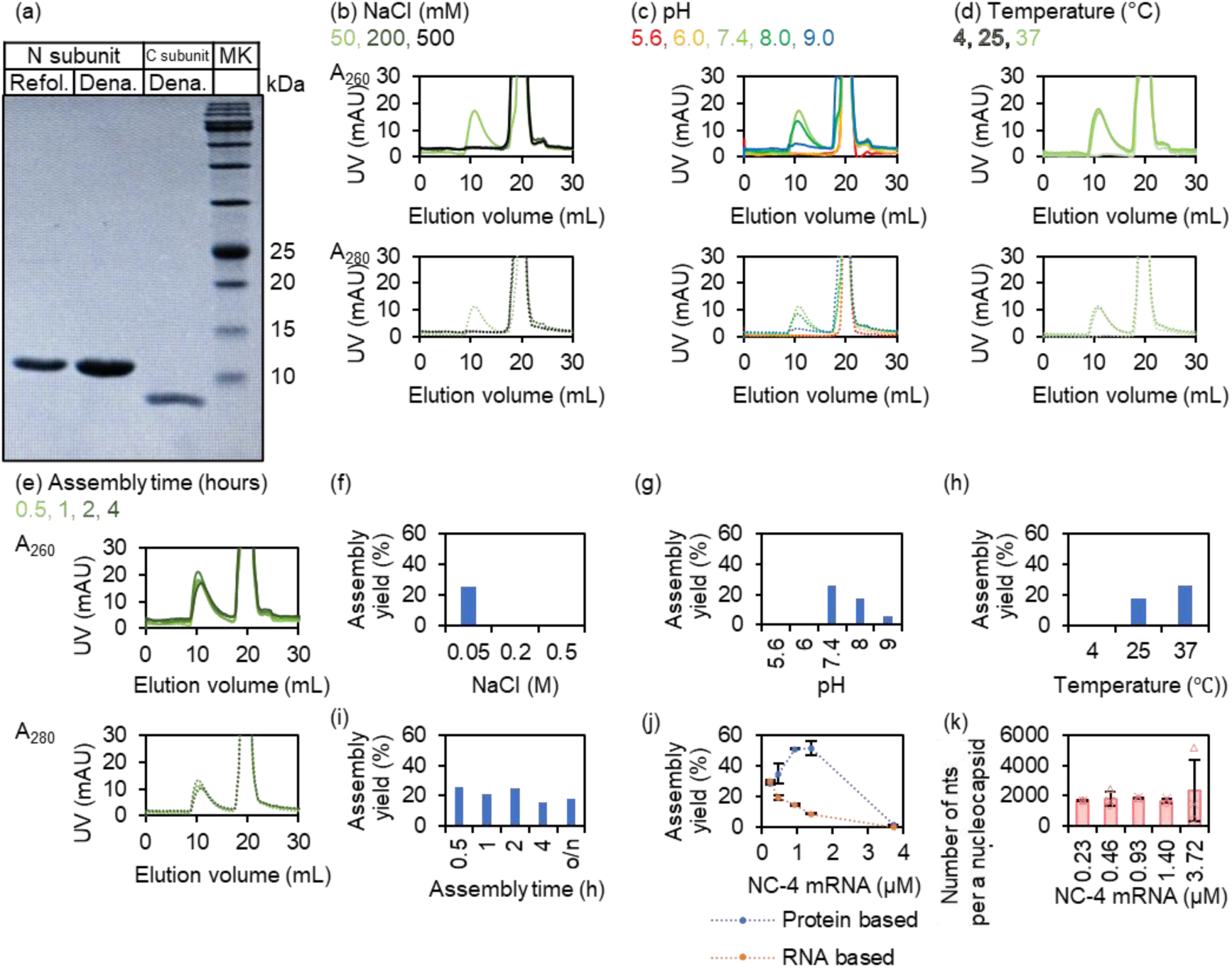
Condition screening of in vitro assembly of split capsid proteins and NC-4 mRNA. **(a)** SDS PAGE analysis of purified N subunit and C subunit. Refol.: refolded, Dena.: denatured. **(b)-(e)** Size-exclusion chromatographs of *in vitro* assembly reactions of N subunit, C subunit and NC-4 mRNA with different (b) NaCl concentration, (c) pH, (d) temperature, and (e) assembly time. Except the particular parameters, assembly conditions were fixed as pH 7.4, 50 mM NaCl, at 37 °C for 30 minutes. **(f)-(i)** Assembly yields of spNC-4 based on protein concentration with different (f) NaCl concentration, (g) pH, (h) temperature for 30 minutes, and (i) assembly time. **(j)** Yields of spNC-4 based on protein and RNA concentration with different NC-4 mRNA concentration. **(k)** Number of RNA nt per a capsid prepared with different NC-4 mRNA concentration. The data in j and k are mean ± s.d. for three independent replicates tested.

**Figure S2.**
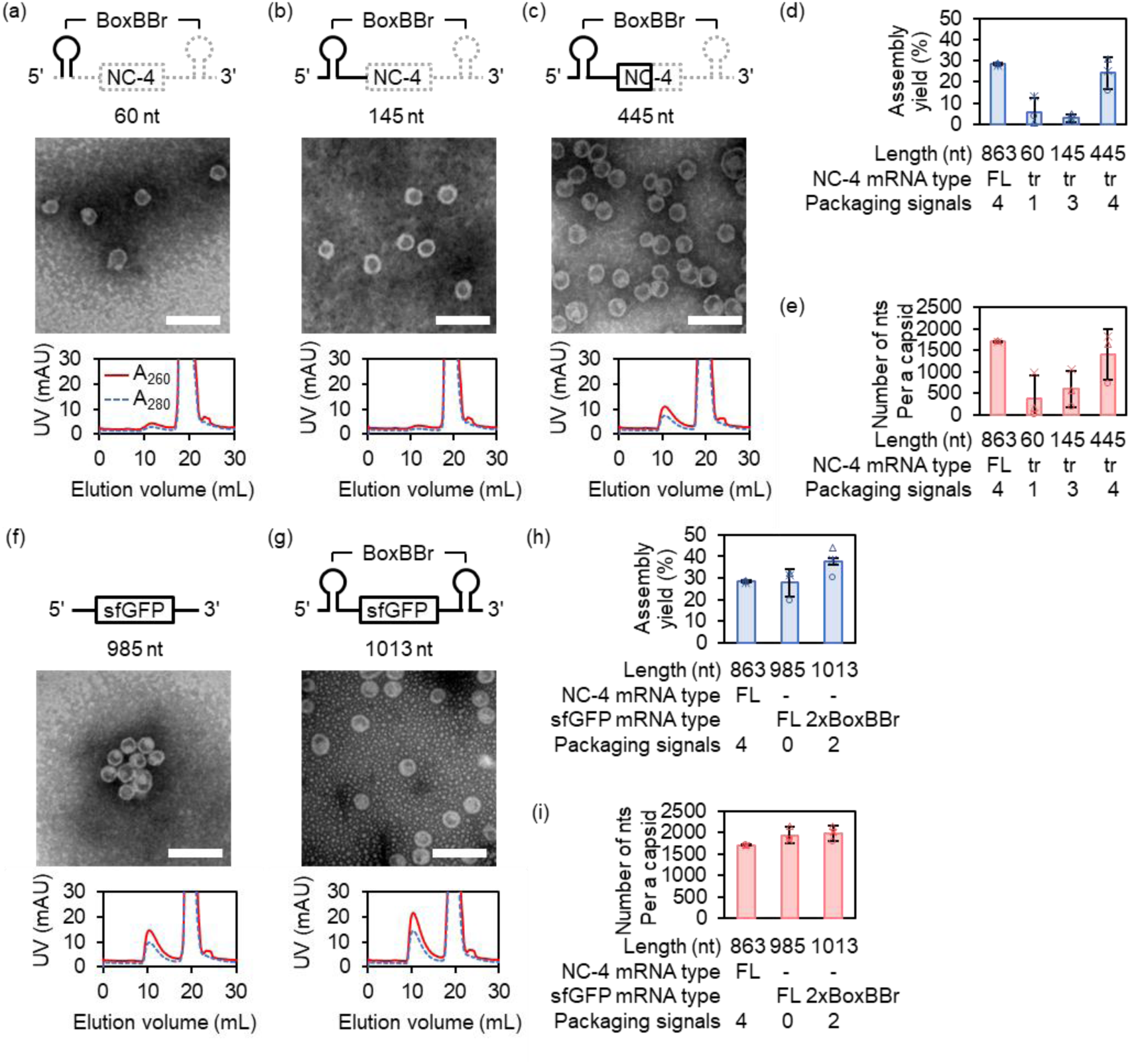
*In vitro* assembly mediated by non-cognate RNAs. **(a)-(c)** Negative-stain TEM analysis and size-exclusion chromatographic (SEC) analysis of *in vitro* assembly mediated by a series of truncated NC-4 mRNAs with 60, 145 or 445 nt-length. 68.2 ng/μL each RNA was used for in vitro assembly, corresponding to the 0.23 μM of NC-4 mRNA with 863 nt-length. Scale bars: 100 nm. **(d)** *In vit*ro assembly yields based on protein concentration, which was mediated by truncated NC-4 mRNAs. FL: full-length, tr: truncated. **(e)** Number of RNA nts per a capsid as a result of *in vitro* assembly mediated by truncated NC-4 mRNAs. FL: full-length, tr: truncated. **(f), (g)** Negative-stain TEM analysis and SEC analysis of *in vitro* assembly mediated by (f) mRNA of sfGFP and (g) mRNA of sfGFP fused with 2x BoxBBr packaging signals. mRNA concentration was 0.23 μM. Scale bars: 100 nm. **(h)** *In vit*ro assembly yields based on protein concentration, which was mediated by NC-4 mRNA, mRNA of sfGFP and mRNA of sfGFP fused with 2x BoxBBr packaging signals. FL: full-length, tr: truncated. **(i)** Number of RNA nts per a capsid as a result of *in vitro* assembly mediated by NC-4 mRNA, mRNA of sfGFP and mRNA of sfGFP fused with 2x BoxBBr packaging signals. FL: full-length, tr: truncated. The data in d, e, h and i are mean ± s.d. for three independent replicates tested.

**Figure S3.**
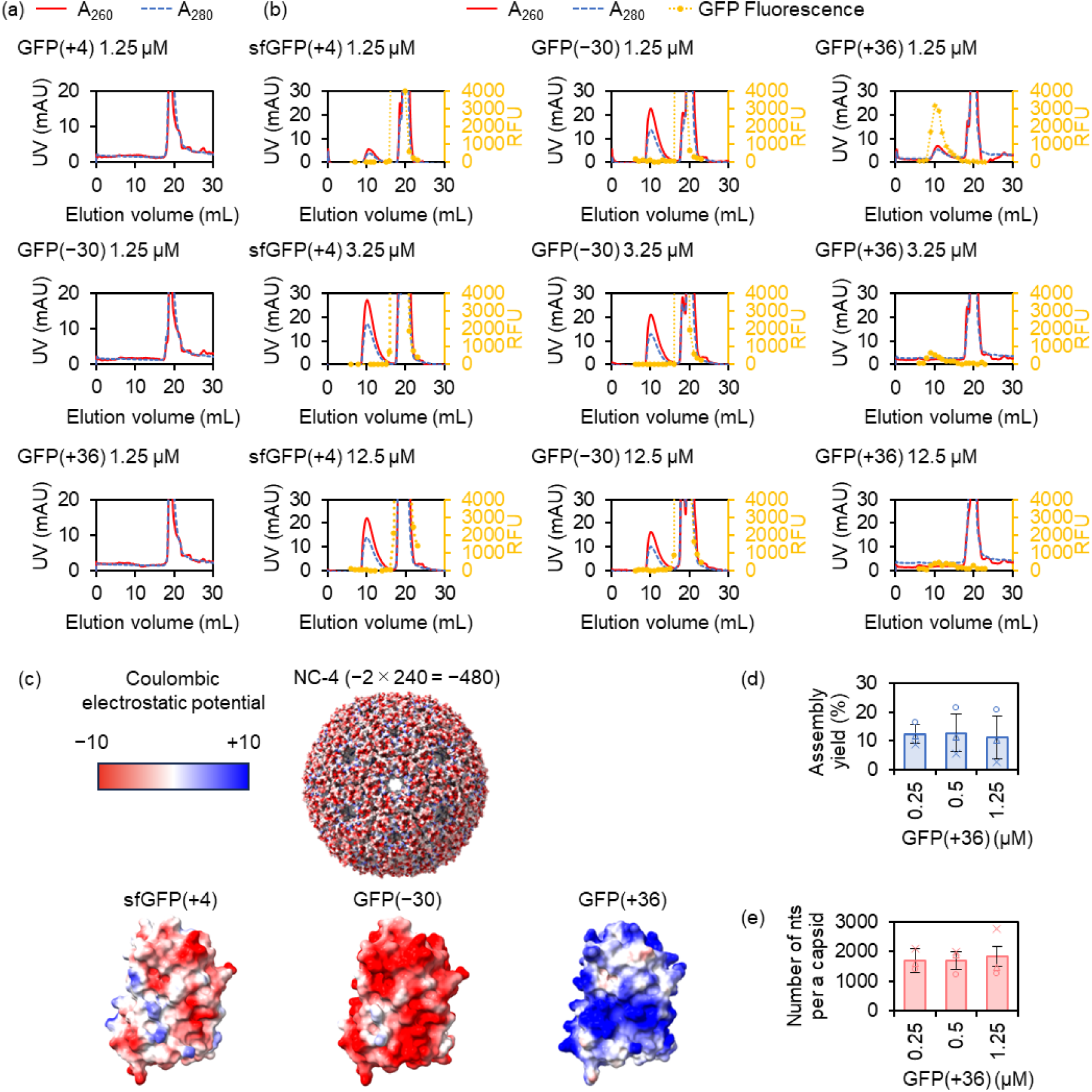
*In vitro* assembly attempts mediated by proteins, and RNA and proteins. **(a)** Size-exclusion chromatographs of i*n vitro* assembly attempts of N subunit, C subunit and GFP(+4), GFP(−30) or GFP(+36). **(b)** Size-exclusion chromatographs of *in vitro* assembly attempt of N subunit, C subunit, NC-4 mRNA and 1.25 μM, 3.75 μM, or 12.5 μM of sfGFP(+4), GFP(−30) or GFP(+36). RFU: relative fluorescence unit. **(c)** Coulombic electrostatic potential on outer surface of NC-4 (PDB ID: 7A4J), GFP(+4) (2B3P), GFP(−30) and GFP(+36). Molecular graphics of GFP(−30) and GFP(+36) were generated based on GFP(+4). **(d)** Yields of nucleocapsid packaging GFP(+36) based on protein concentration assembled *in vitro* with different GFP(+36) concentration. **(e)** Number of RNA nts per a capsid prepared assembled *in vitro* with different GFP(+36) concentration. The data in d and e are mean ± s.d. for three independent replicates tested.

**Figure S4.**
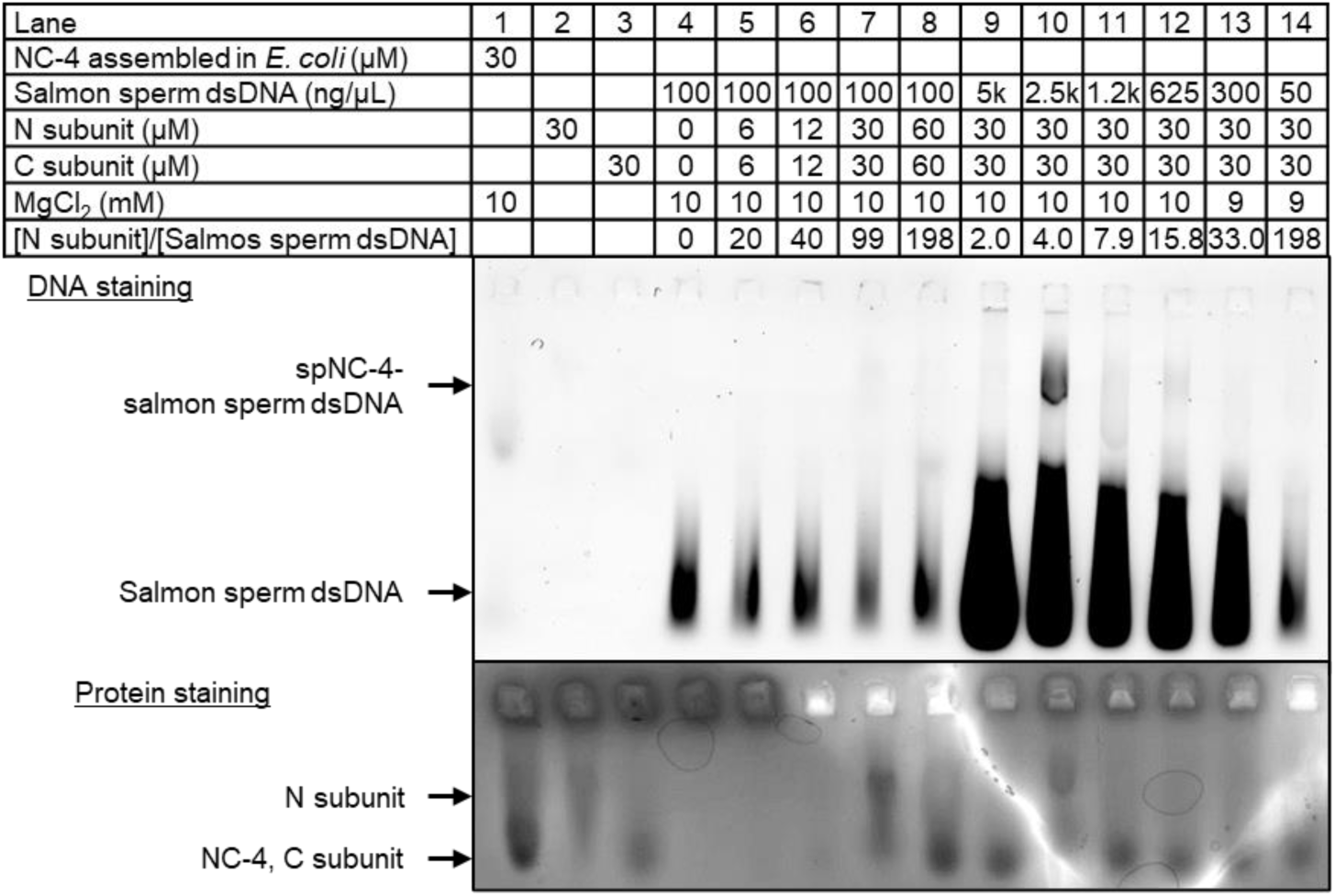
*In vitro* assembly of salmon sperm dsDNA. 0.6% agarose gel electrophoretic (AGE) analysis of i*n vitro* assembly reaction of N subunit, C subunit, and salmon sperm dsDNA with various protein-to-DNA ratio.

**Figure S5.**
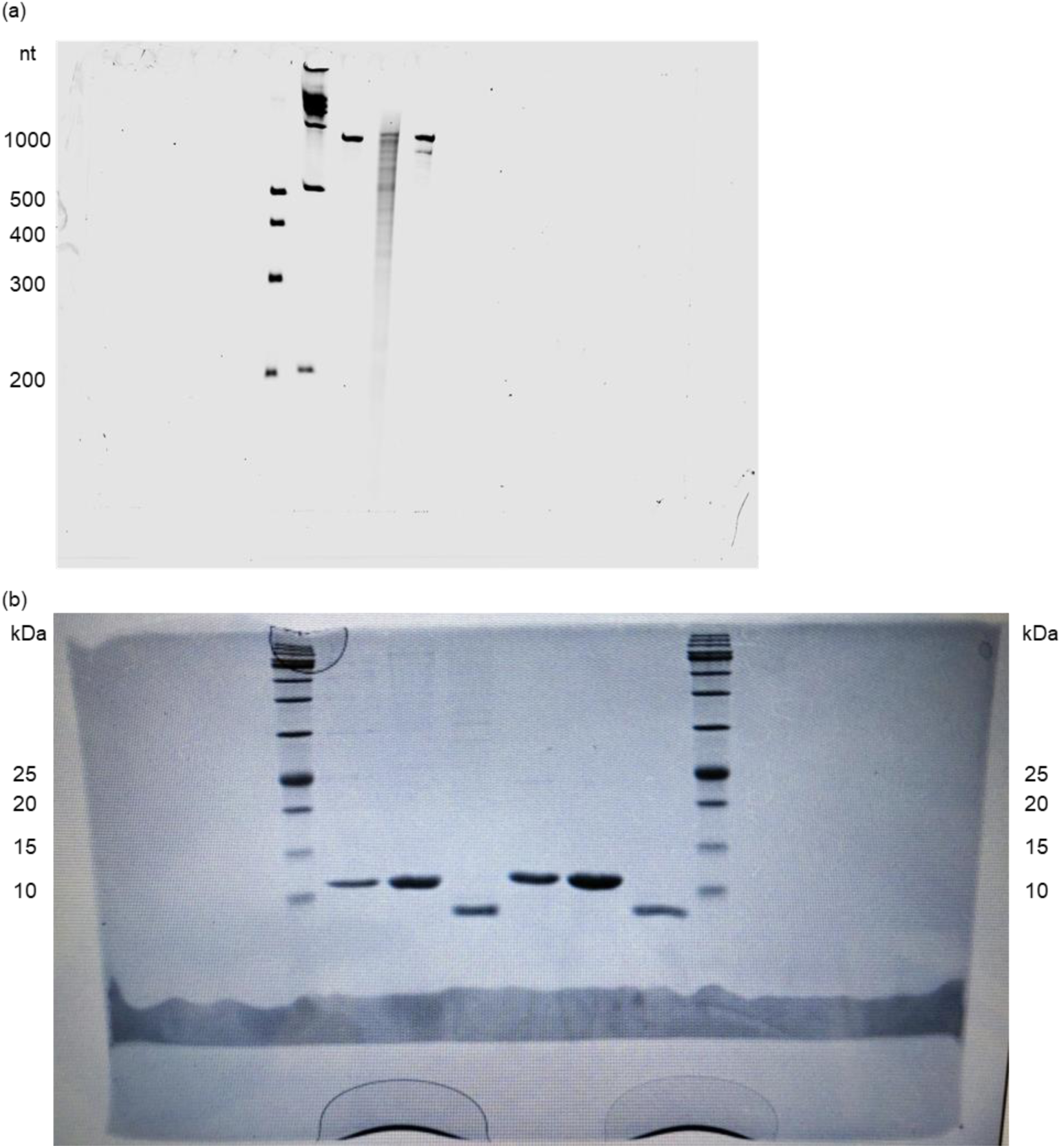
Uncropped gel images **(a), (b)** Uncropped gel images of **(a)** Figure 1d and **(b)** Figure S1a.

**Figure S6.**
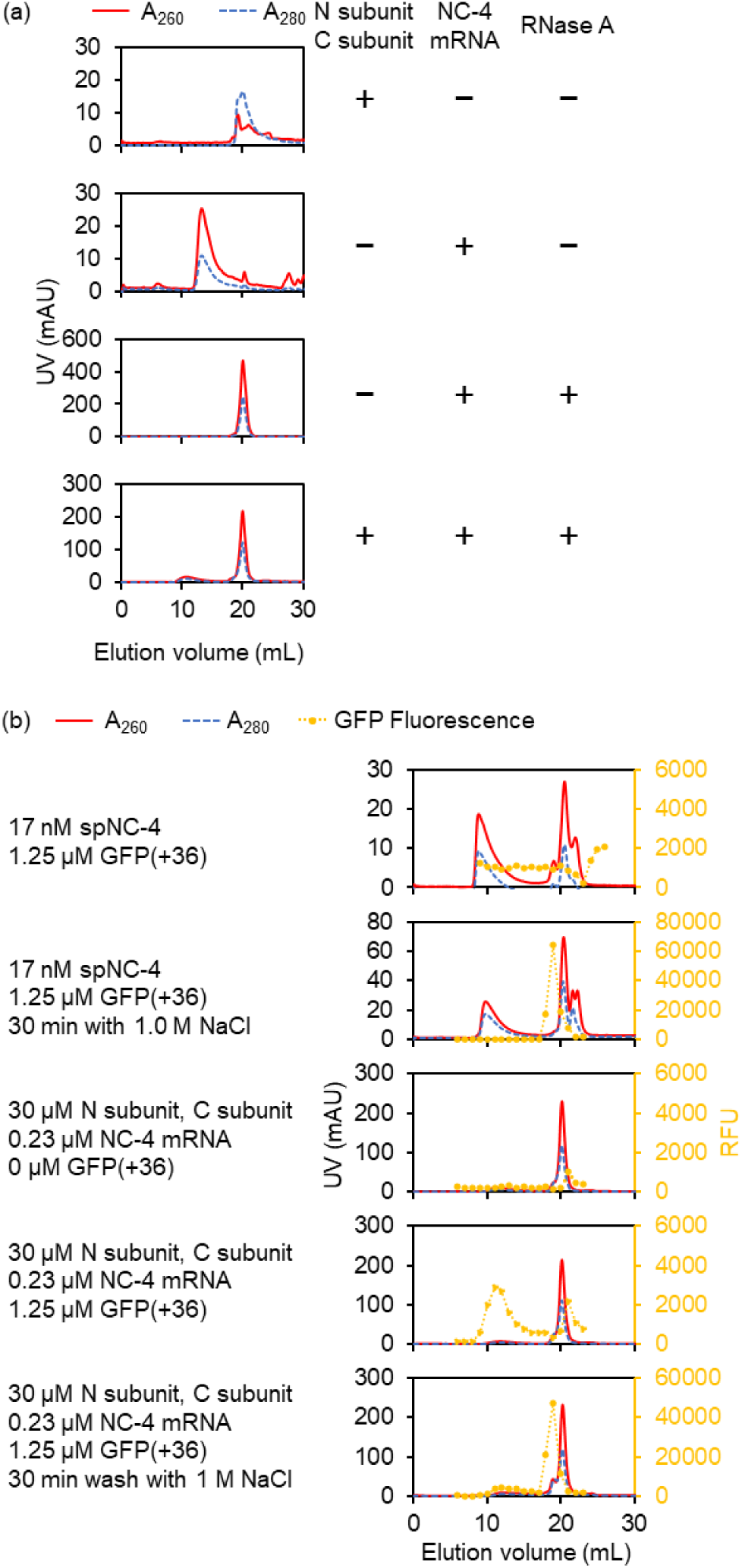
Uncropped SEC chromatographs in Figures **(a), (b)** Uncropped SEC chromatographs in **(a)** Figure 1b, and **(b)** Figure 2b

**Figure S7.**
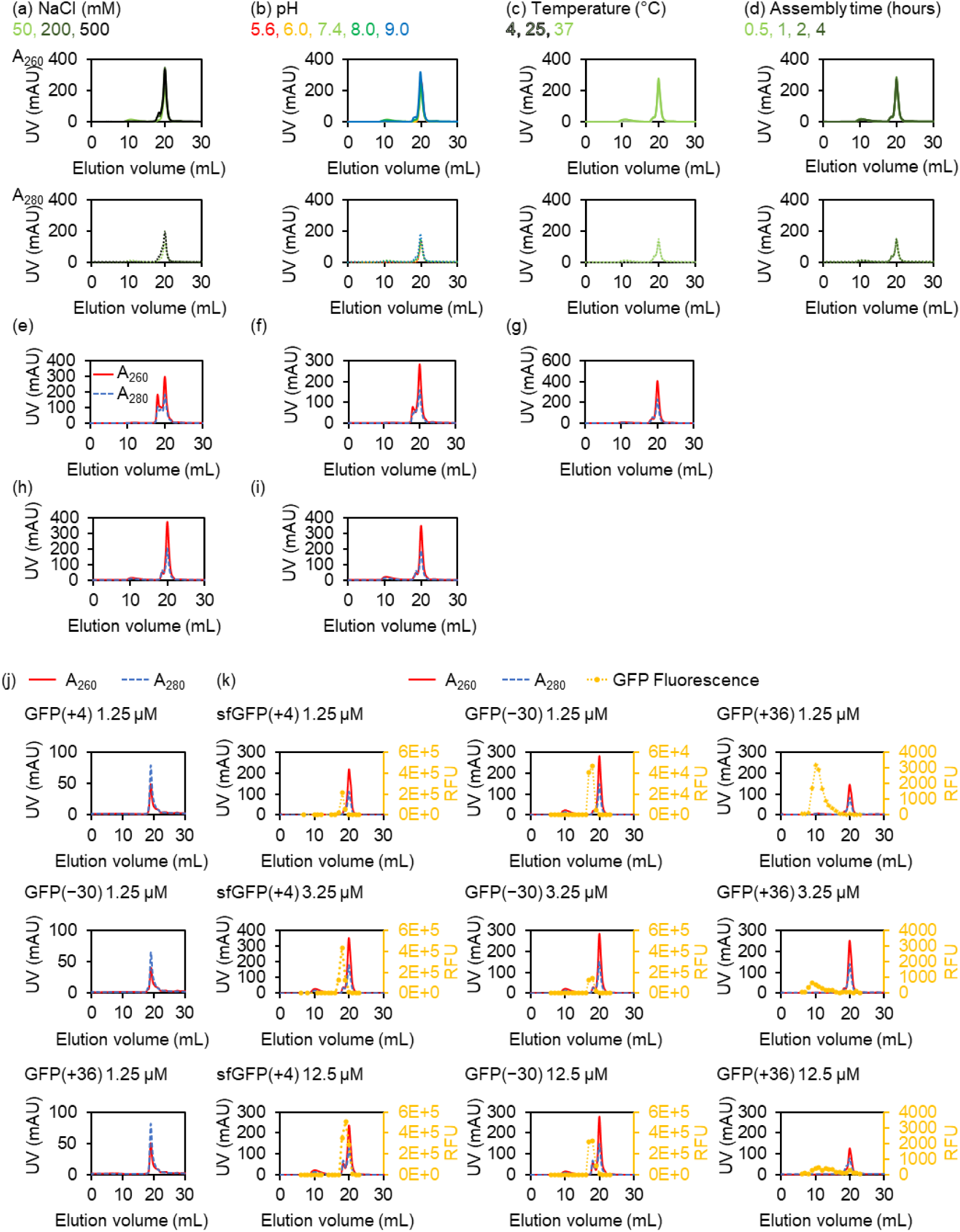
Uncropped SEC chromatographs in Supplementary Figures **(a)-(k)** Uncropped SEC chromatographs in **(a)** Figure S1b, **(b)** Figure S1c, **(c)** Figure S1d, **(d)** Figure S1e, **(e)** Figure S2a, **(f)** Figure S2b. **(g)** Figure S2c, **(h)** Figure S2f, **(i)** Figure S2g, **(j)** Figure S3a and **(k)** Figure S3b.

**Table S1.**
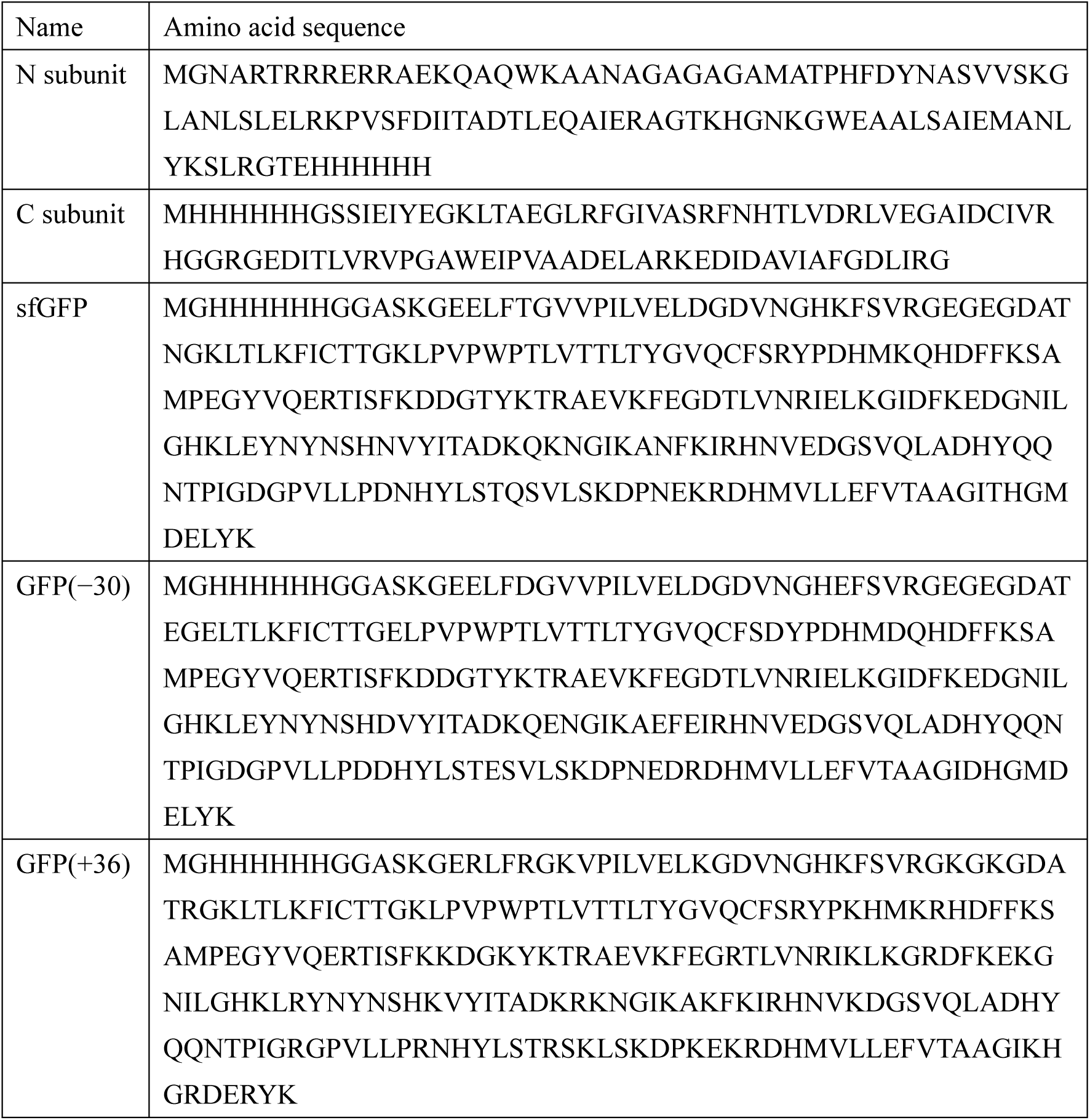
Protein sequences.

**Table S2.**
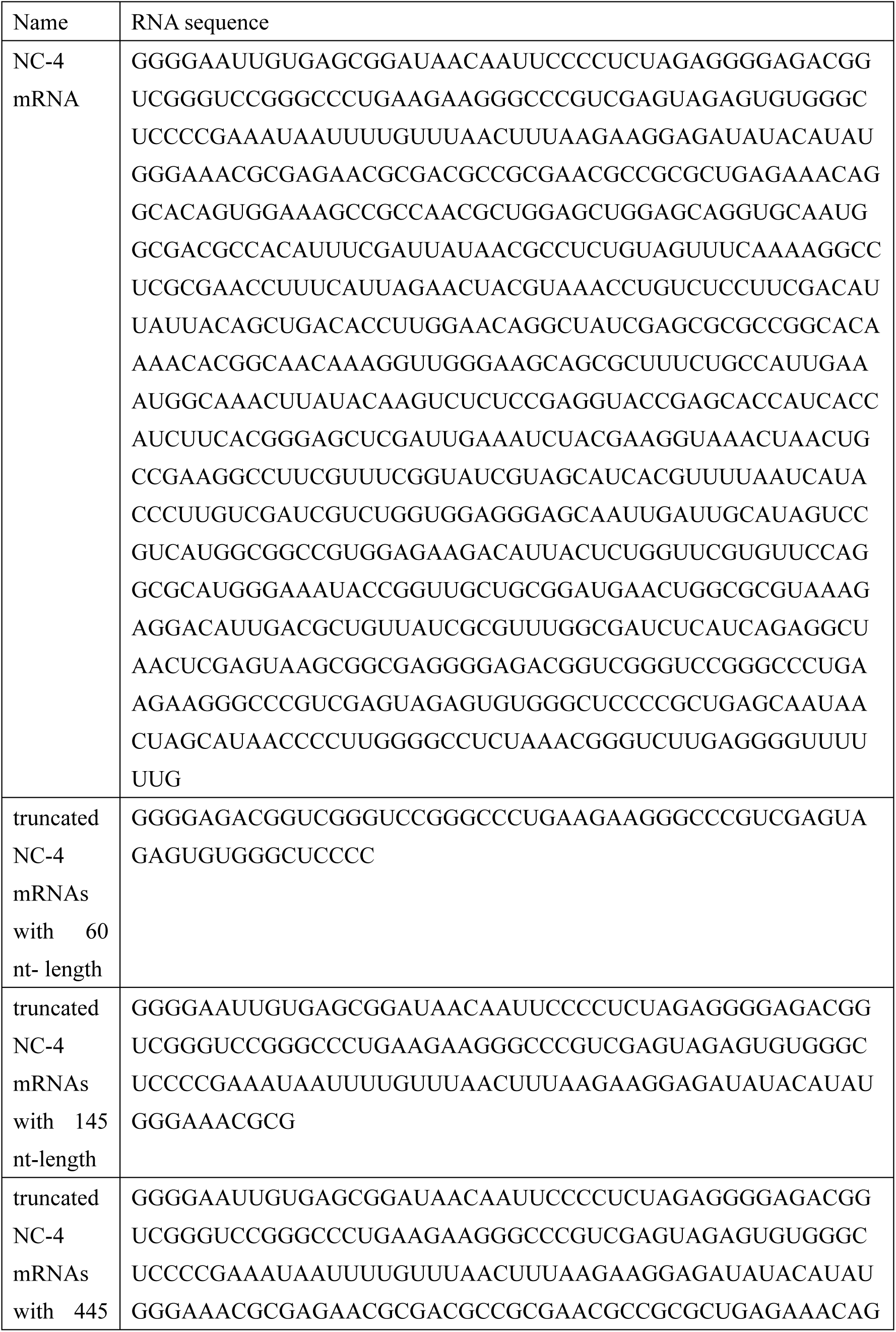
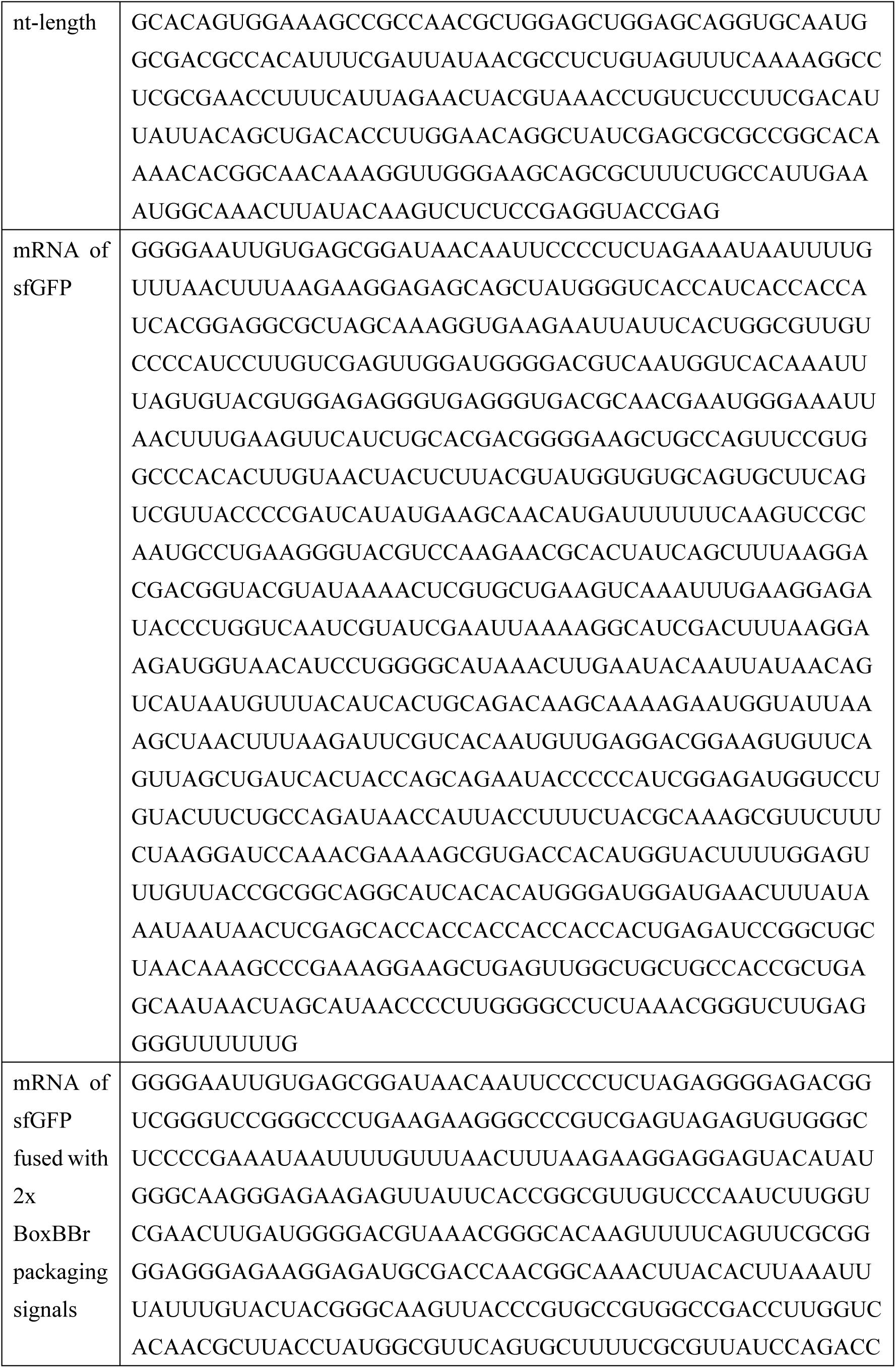
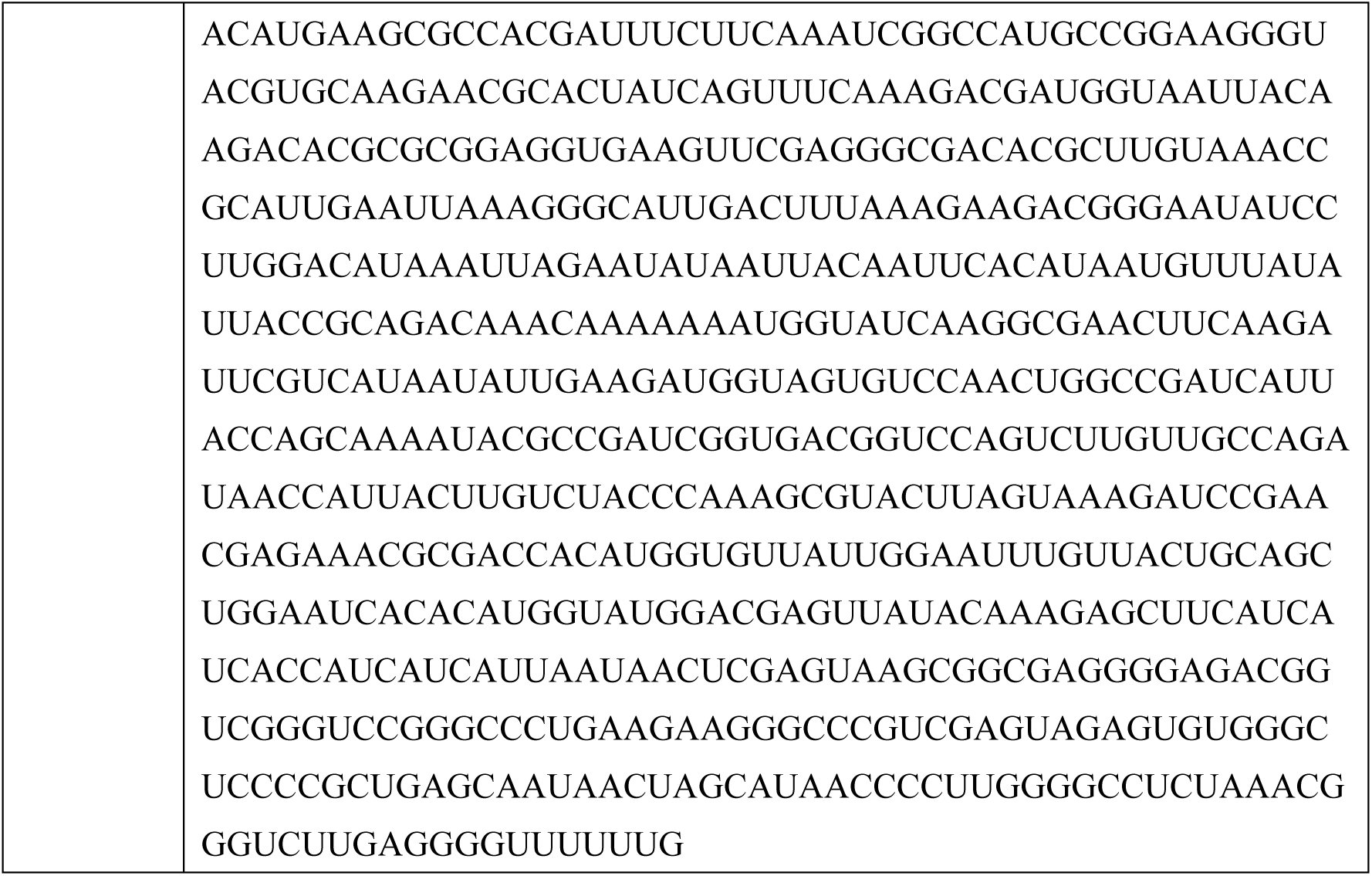
RNA sequences.

**Table S3.**
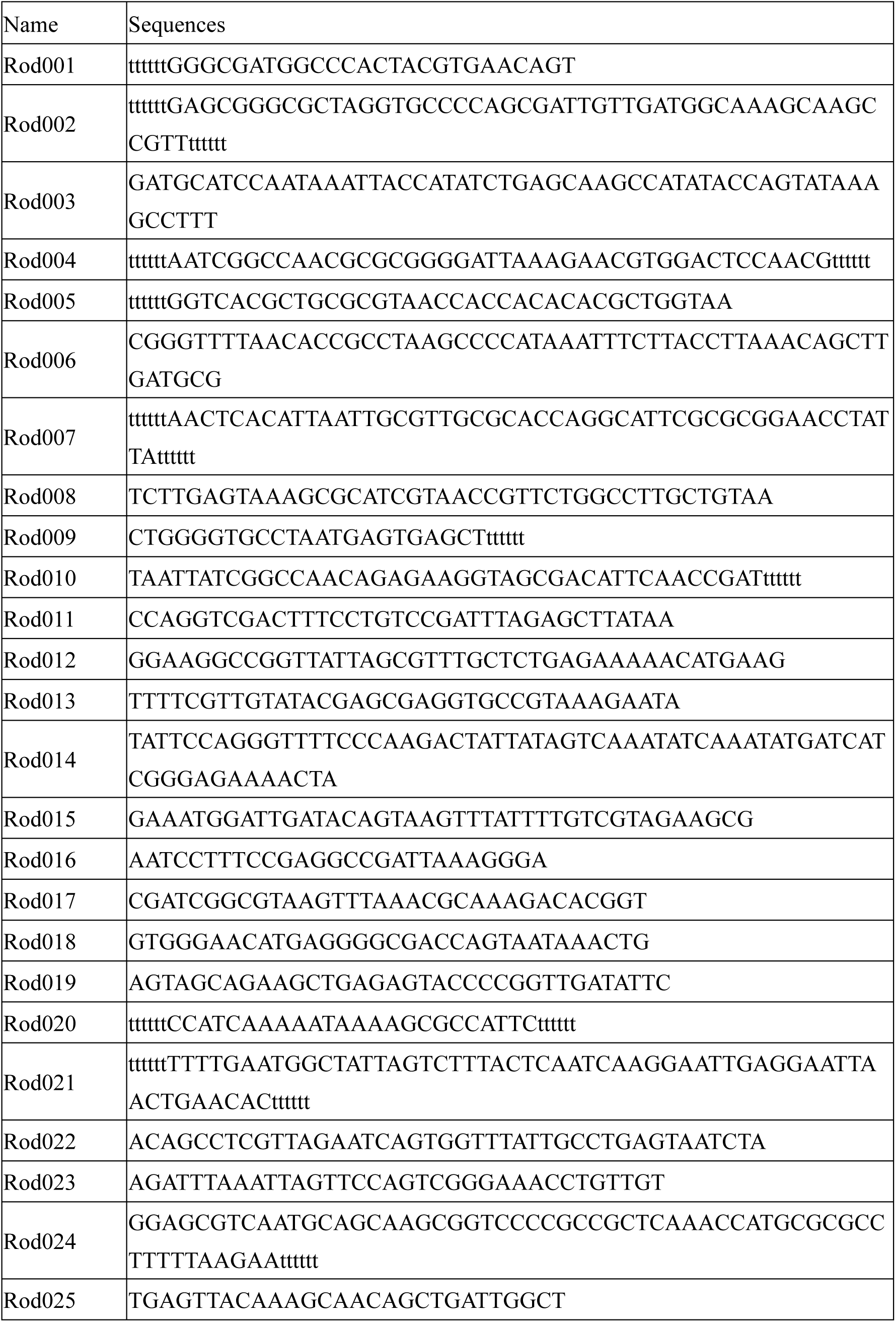
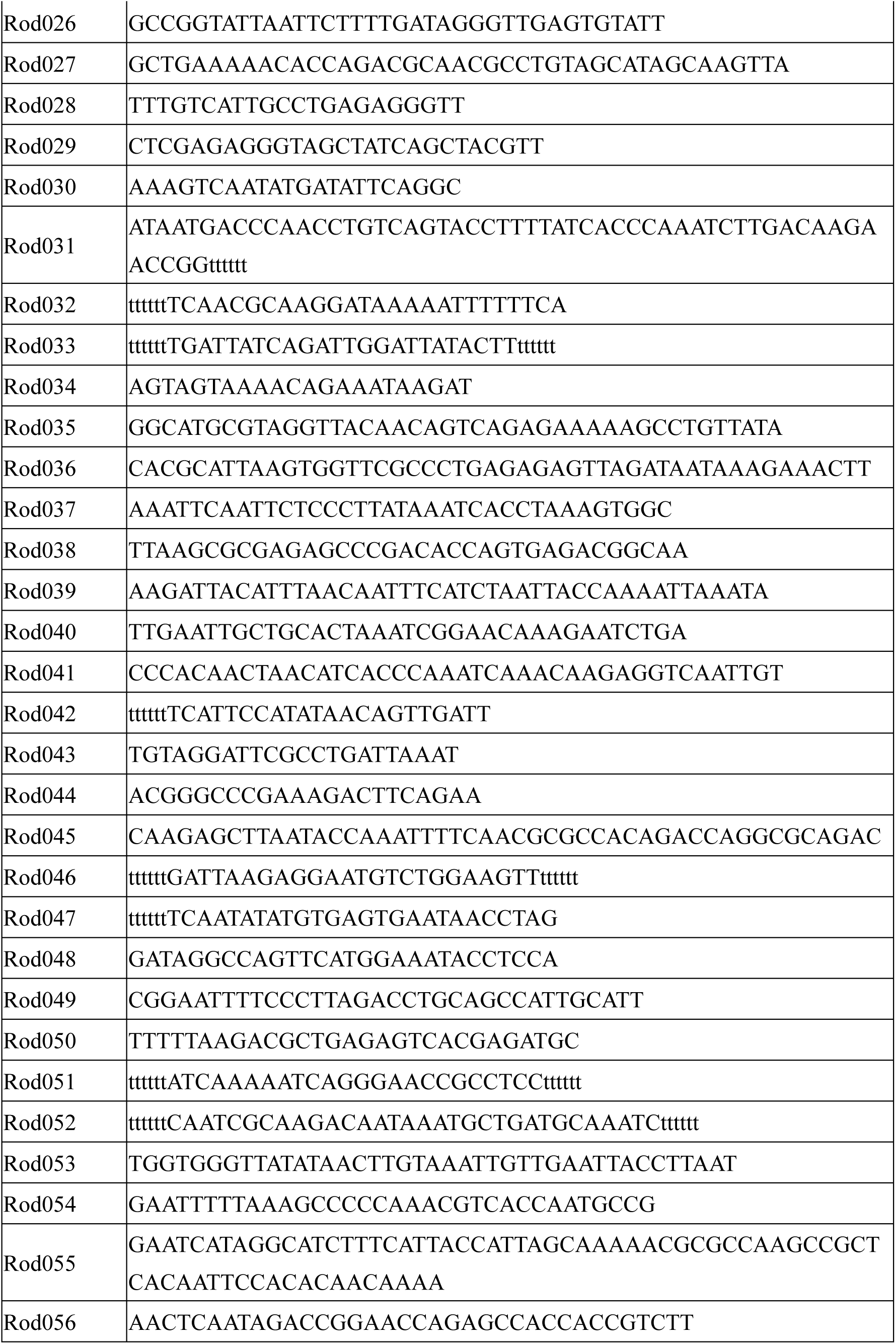
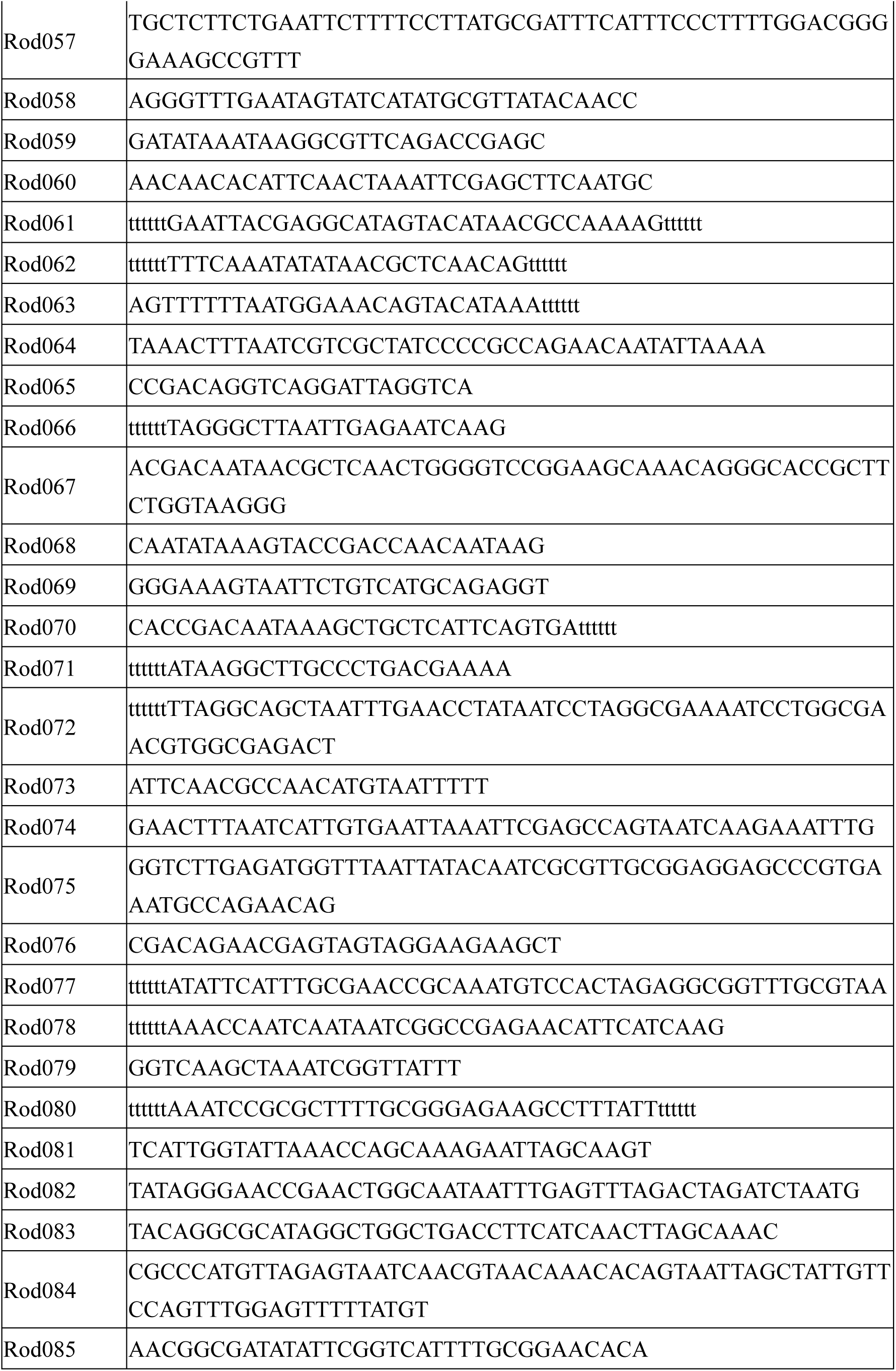
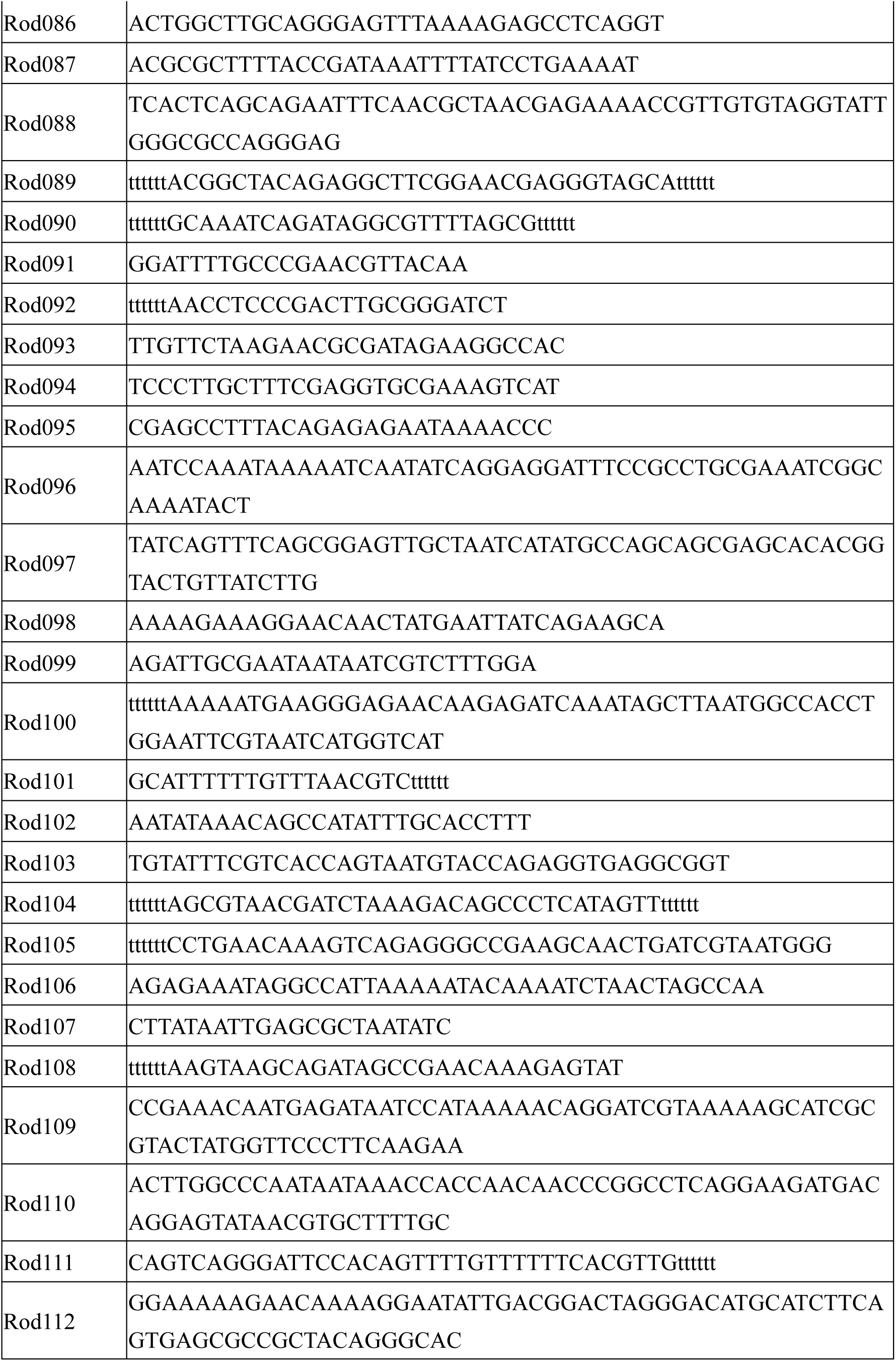
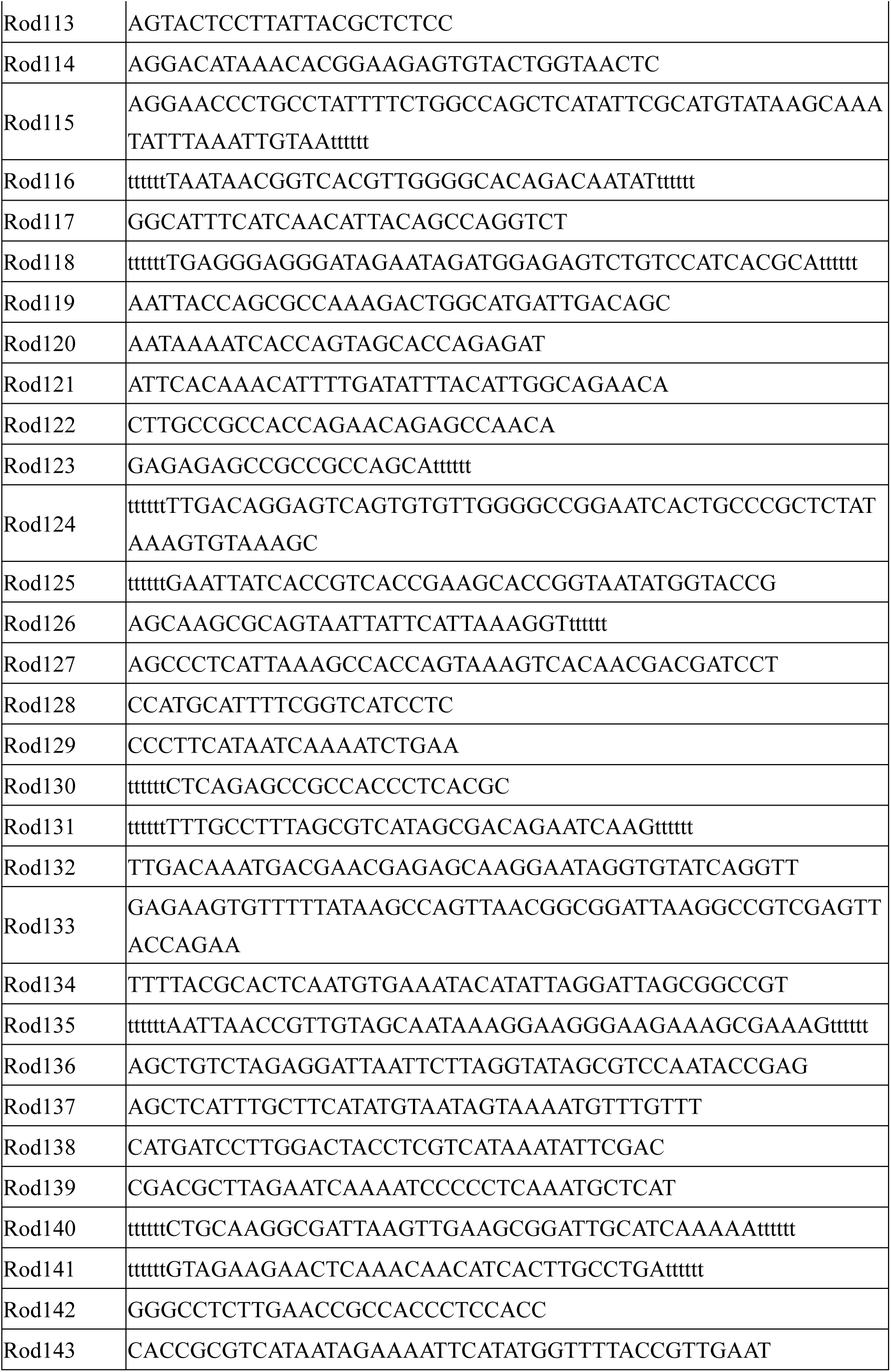
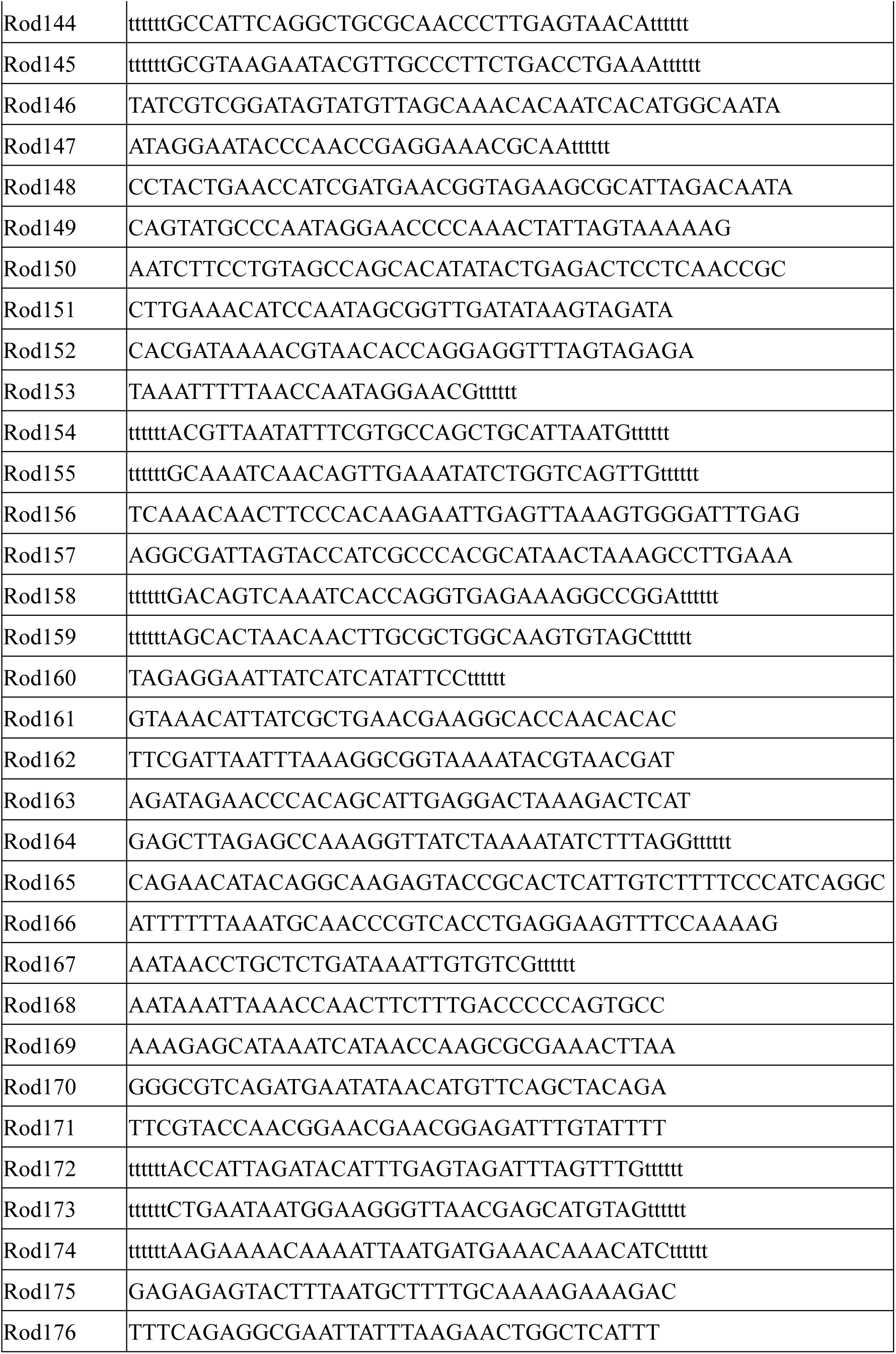
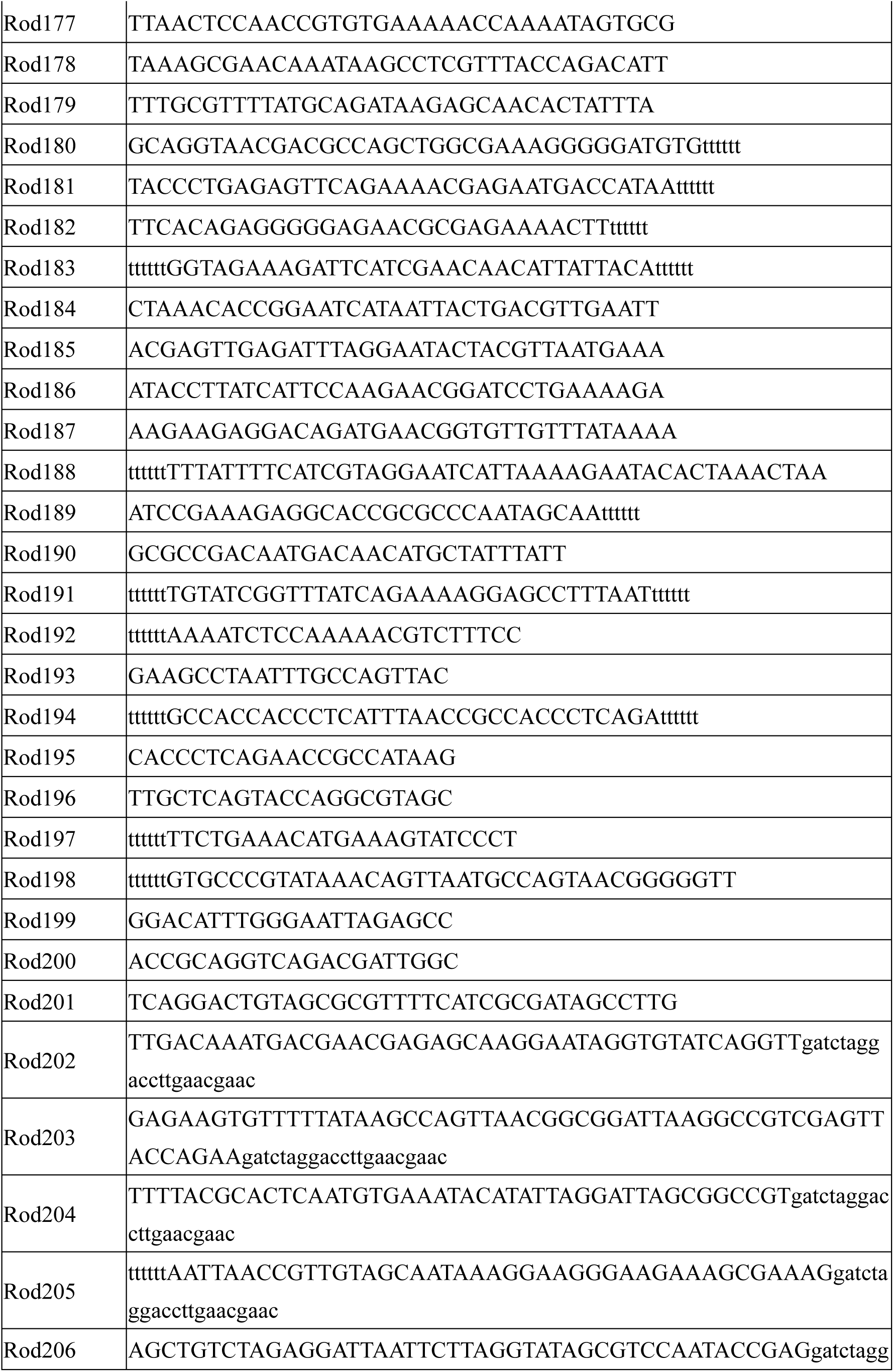
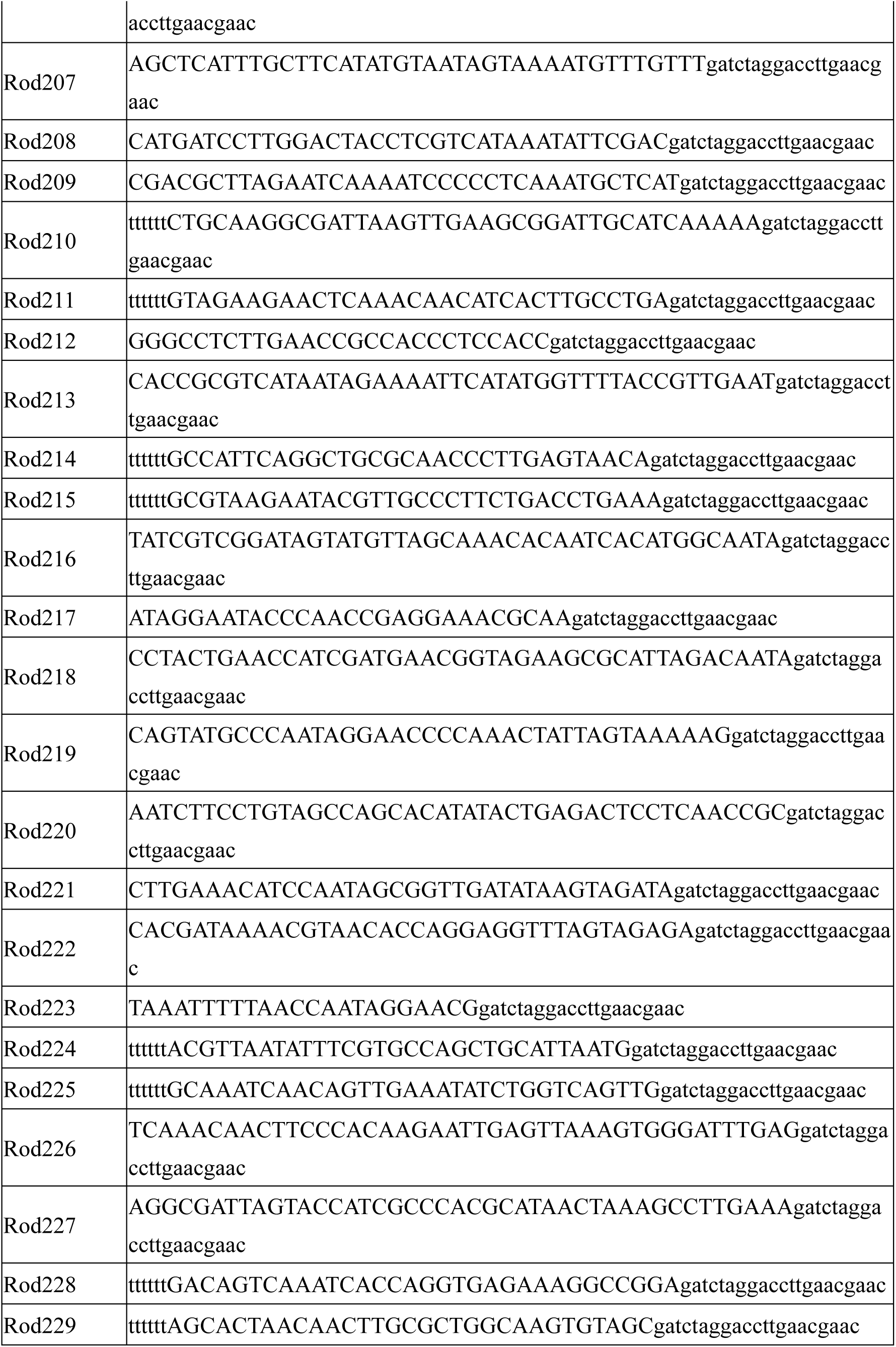
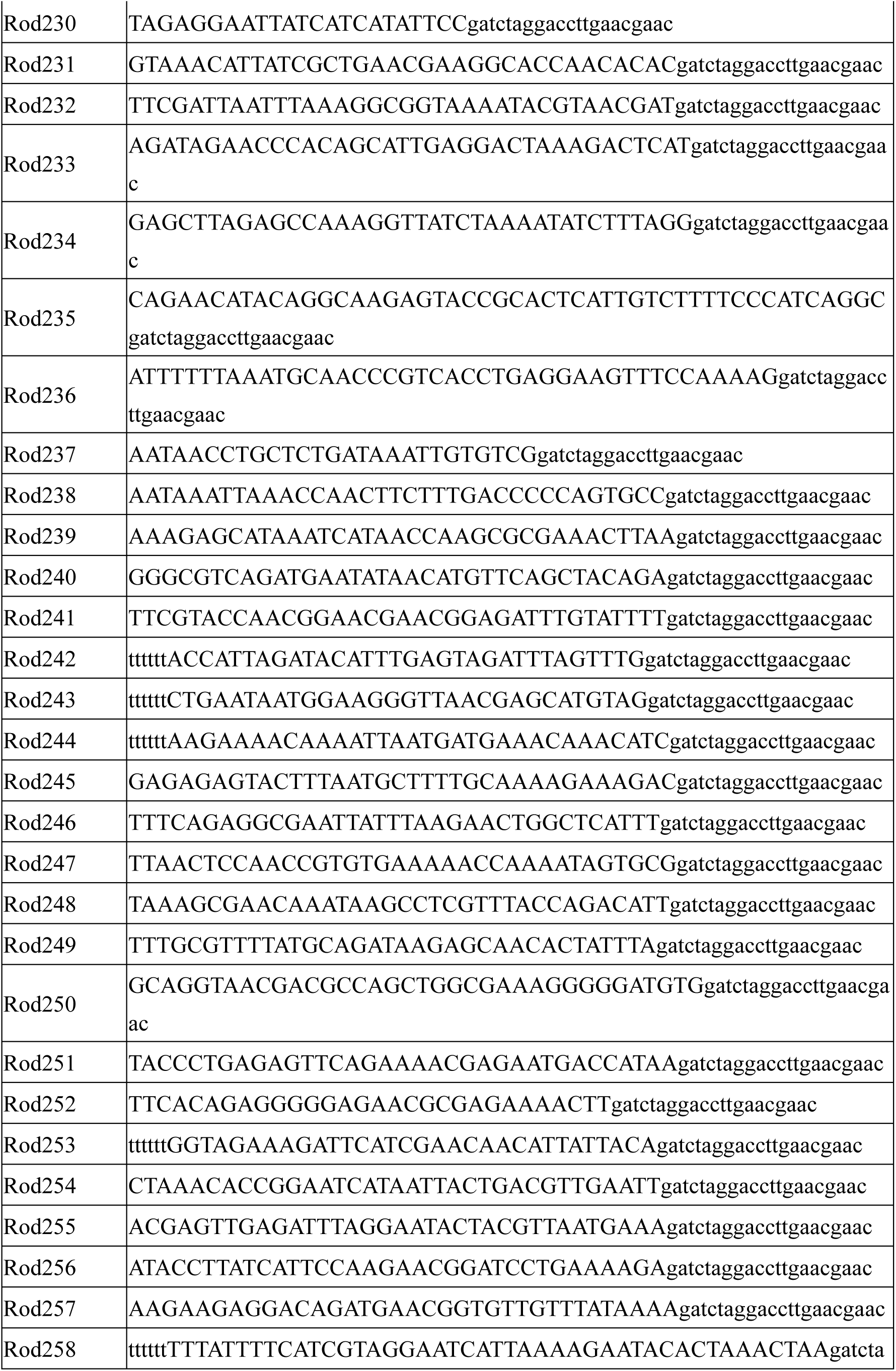
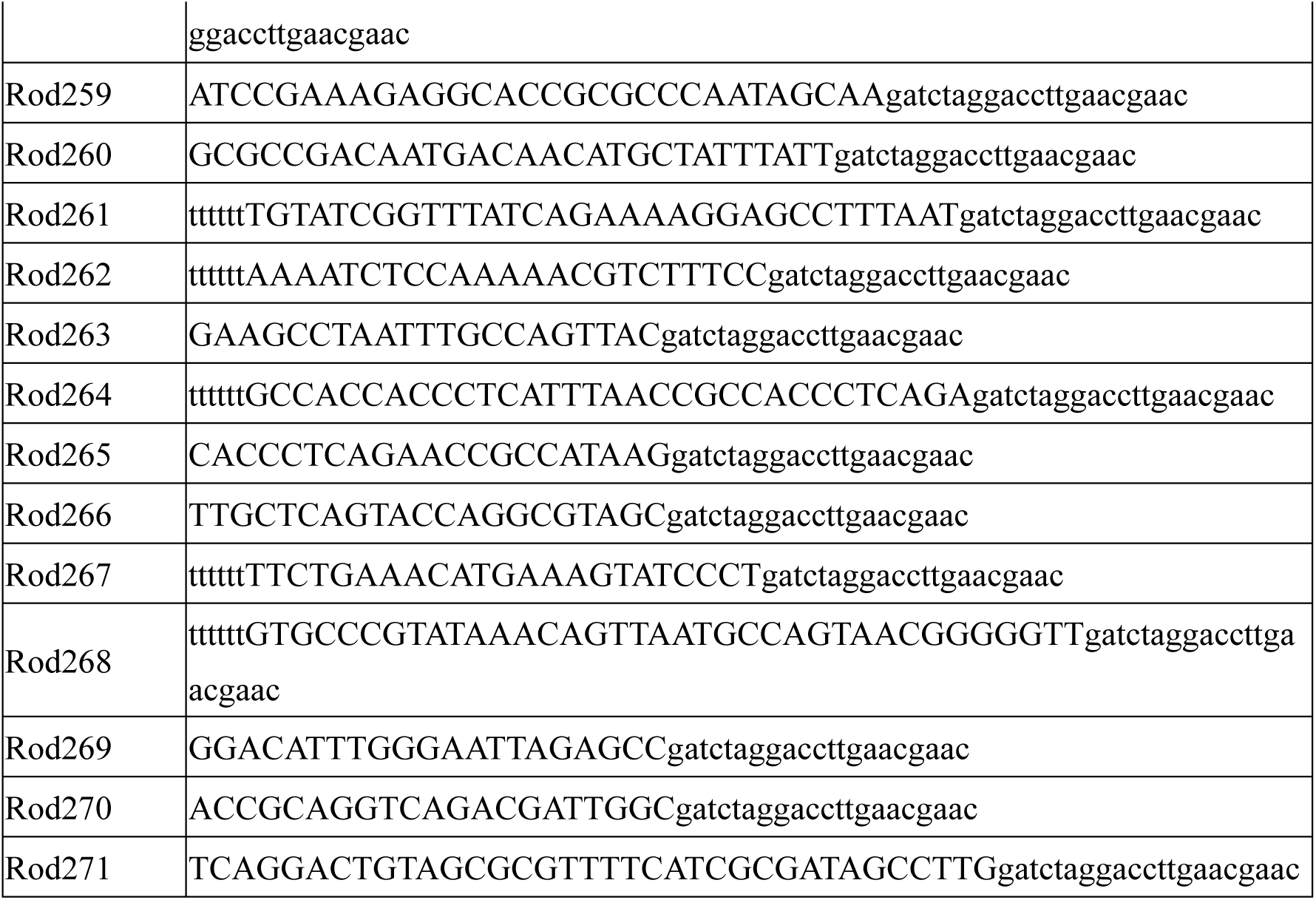
DNA sequences used for DNA origami rod preparation.

**Table S4.**
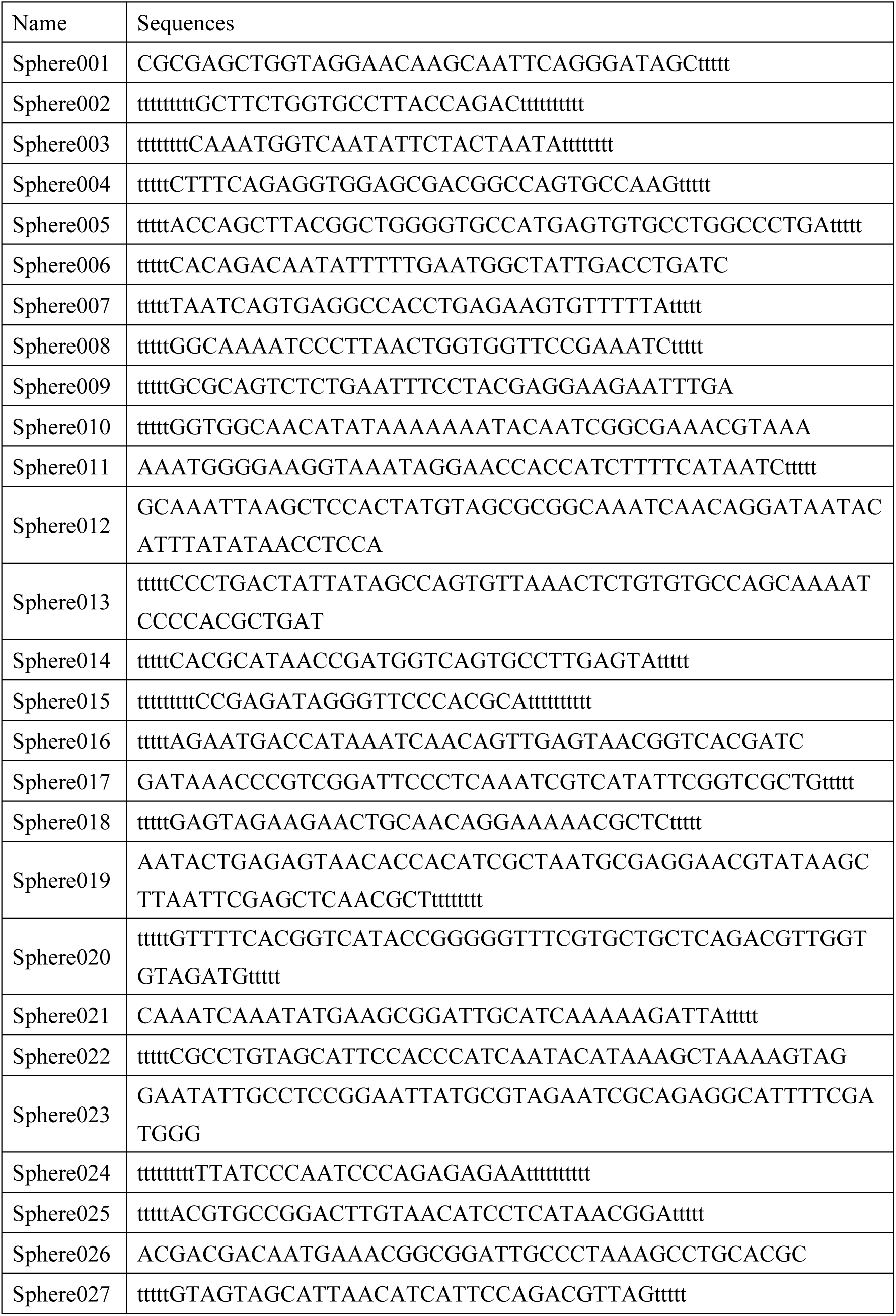
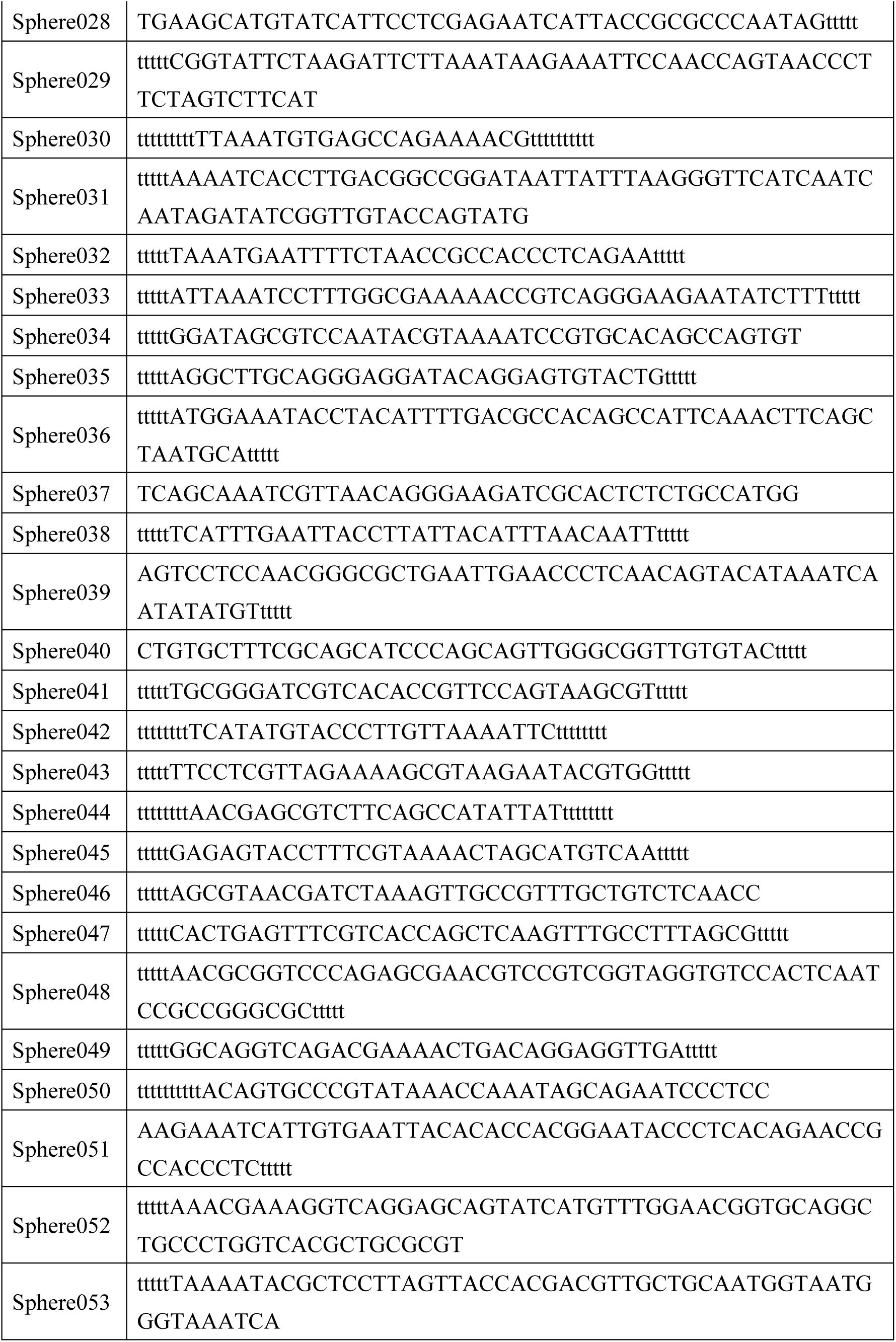
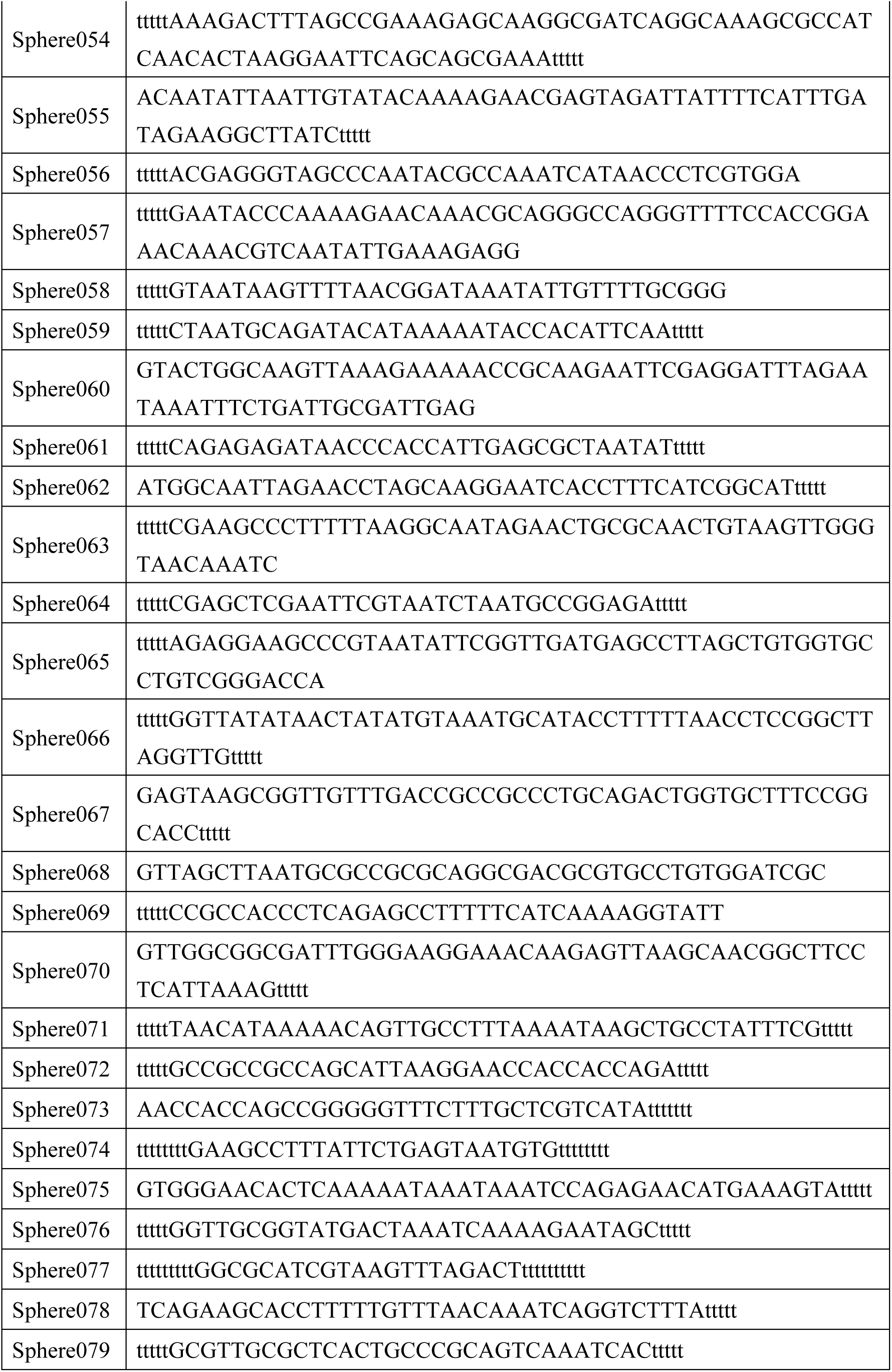
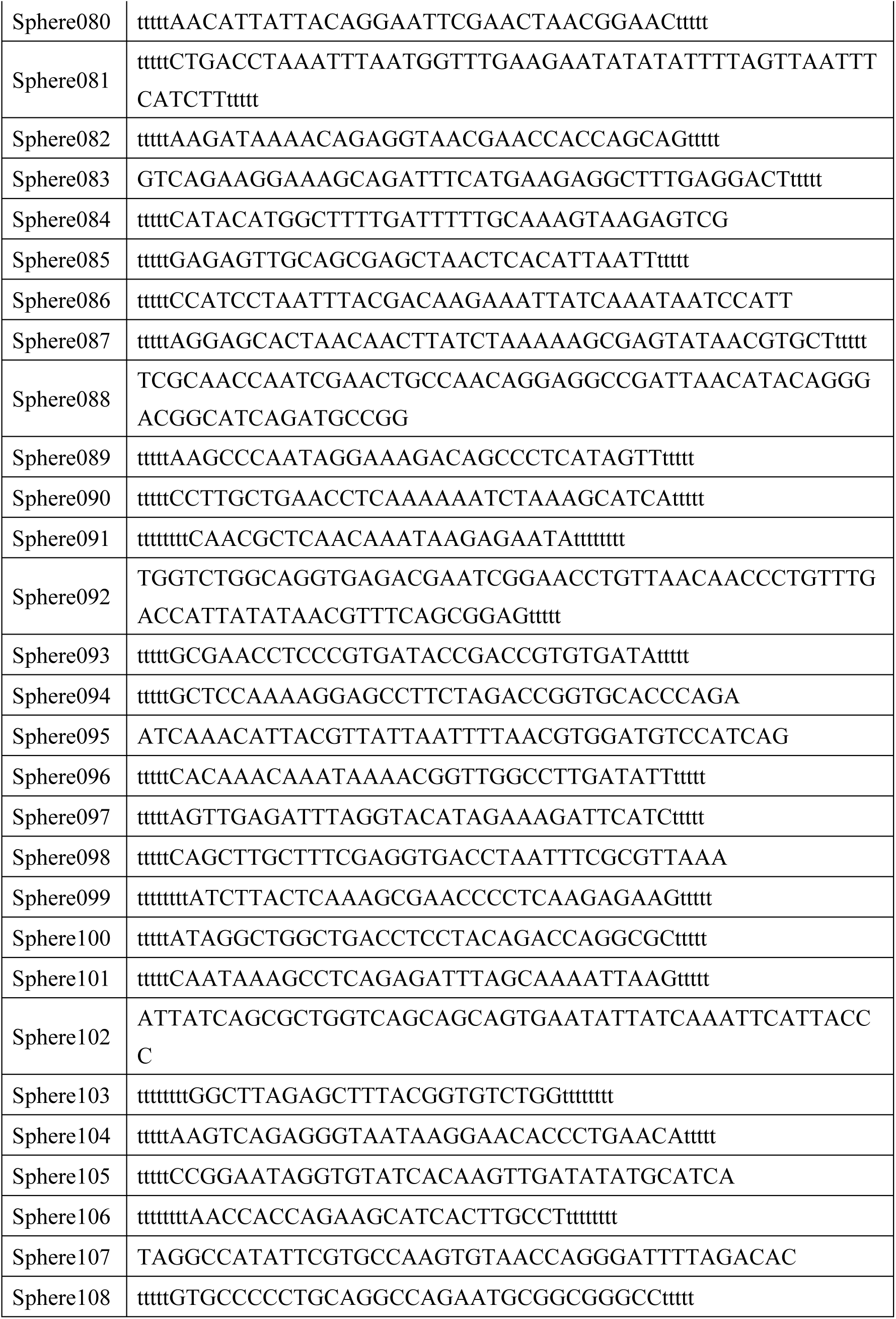
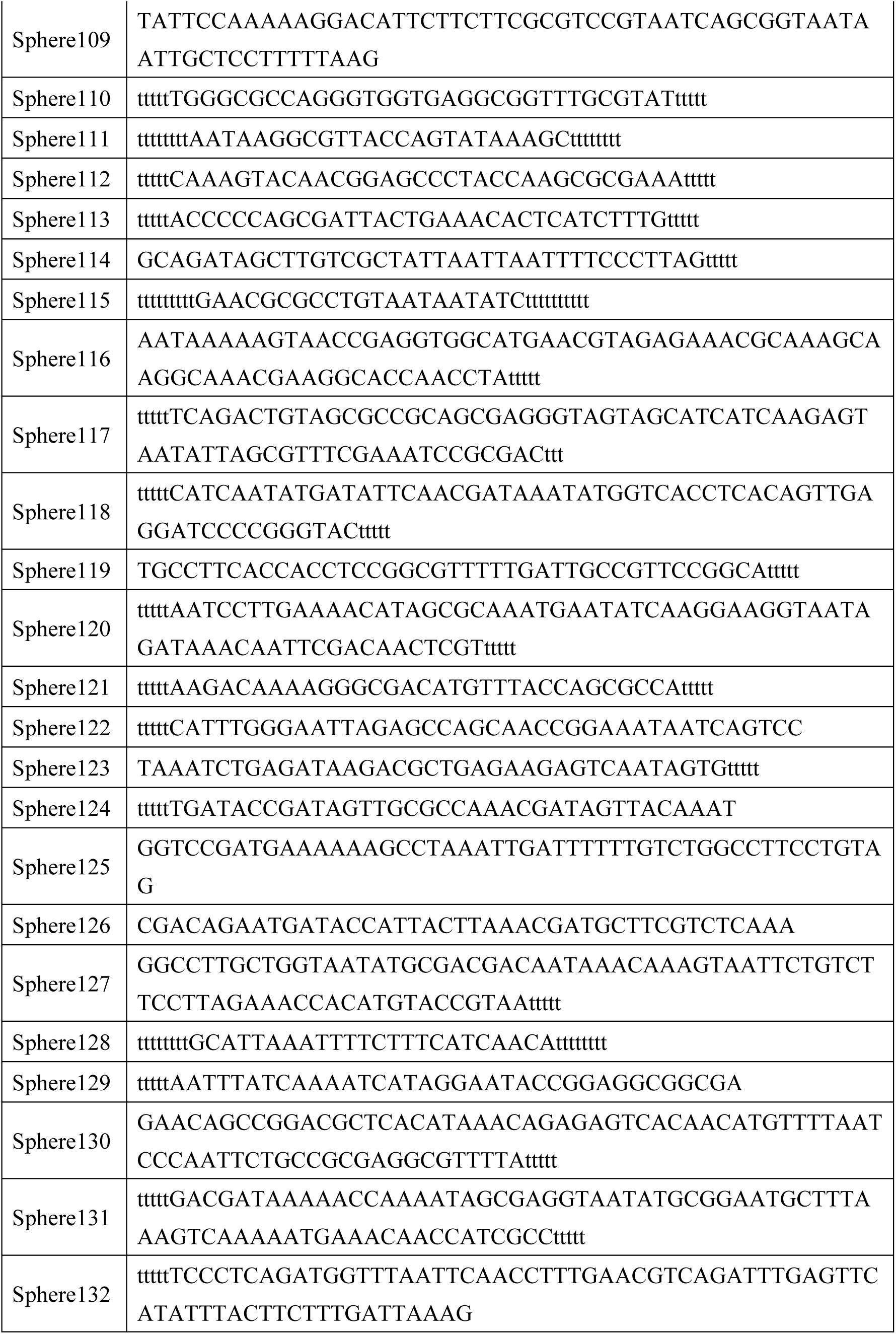
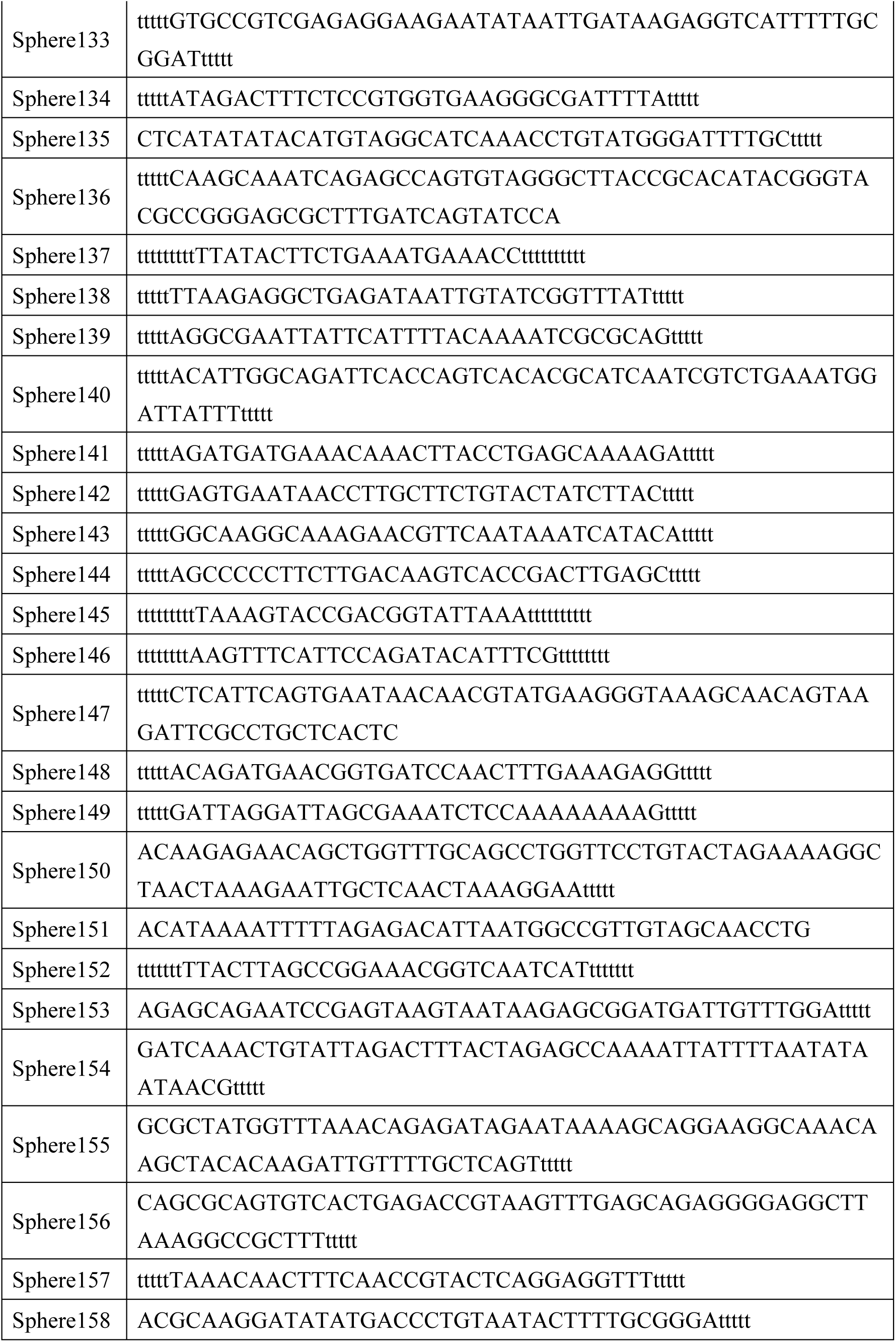
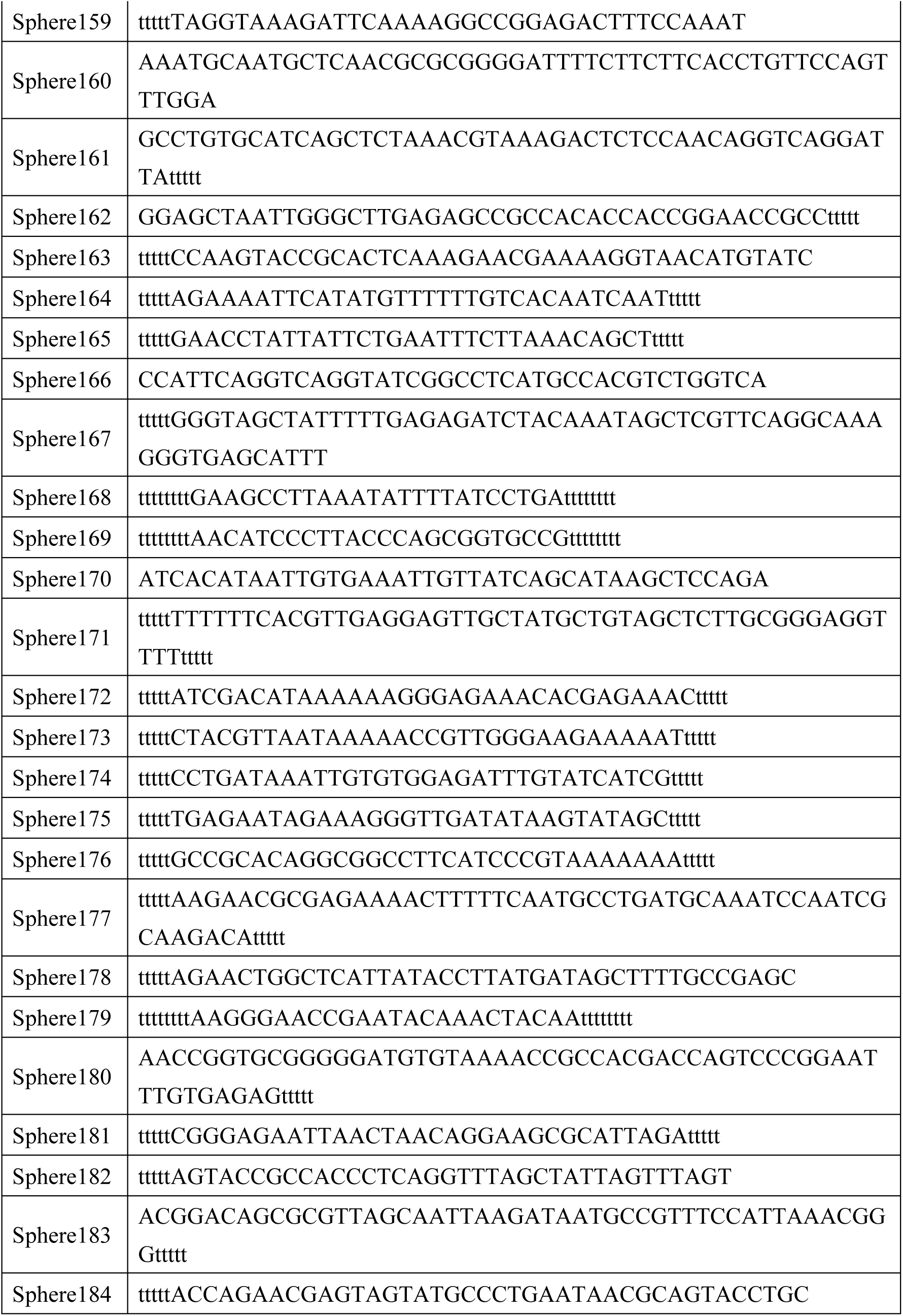
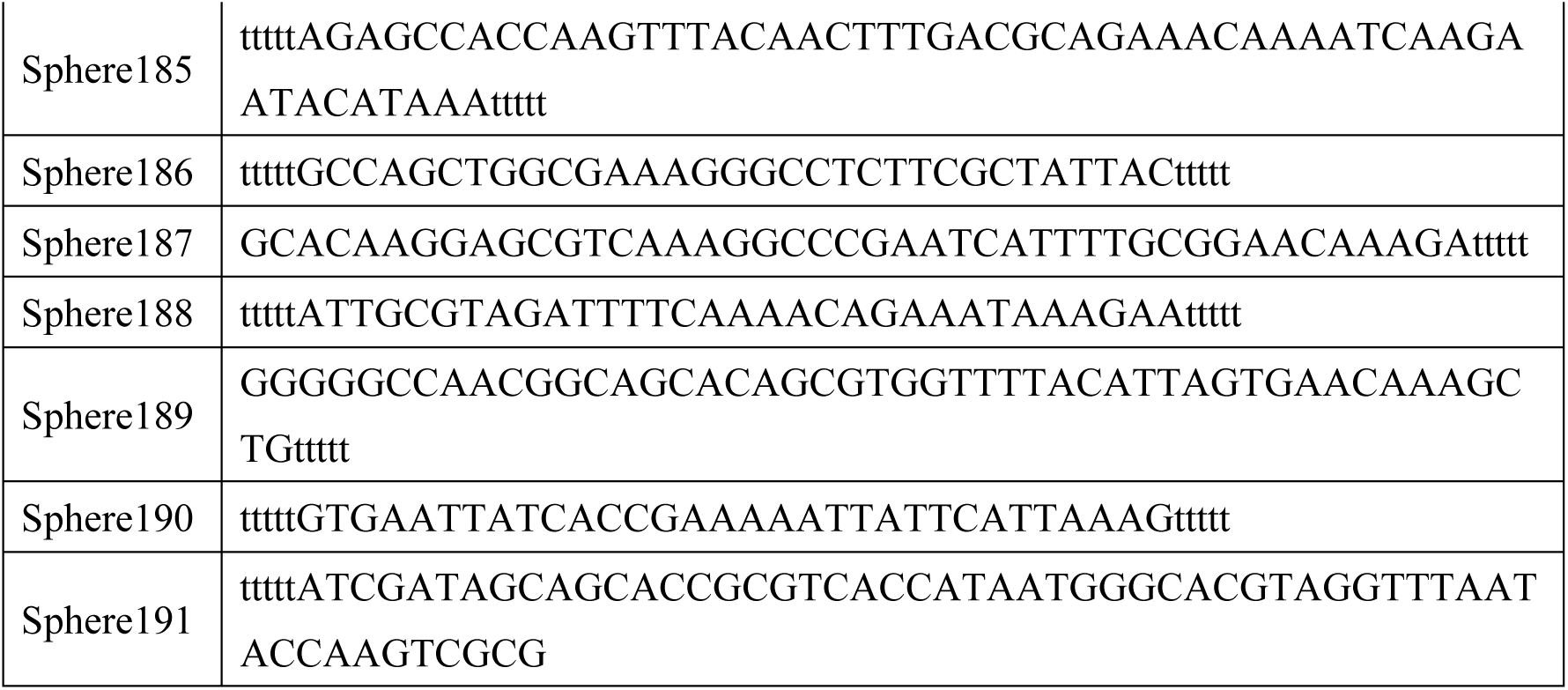
DNA sequences used for DNA origami sphere preparation.

